# Multifaceted brain representation of numerosity across the senses and presentation formats

**DOI:** 10.1101/2025.05.29.653470

**Authors:** Ying Yang, Michele Fornaciai, Irene Togoli, Iqra Shahzad, Remi Gau, Alice Van Audenhaege, Filippo Cerpelloni, Olivier Collignon

## Abstract

Humans can extract numerosity from different senses and a variety of context. How and where the brain abstract numerical information from low-level sensory inputs remains debated. Using multivariate pattern decoding and representational similarity analysis applied to fMRI data, we comprehensively investigate how the brain represents numerical information (range 2-5) across different modalities (auditory, visual) and presentation formats (sequential, simultaneous; symbolic, non-symbolic). We identify a set of brain regions along the dorsal pathway-from early visual cortex to the intraparietal and frontal regions-that encode specific non-symbolic numerical information across formats and modalities. The numerical distance effect, a hallmark of magnitude encoding, was observed in most of these regions. We found aligned representation of numerical information across visual and auditory modalities in intraparietal and frontal regions, but only when they shared a sequential presentation format. Maintaining a distinction between spatial and temporal numerical representations may thus be a fundamental aspect of numerical processing. RSA further revealed a posterior-to-anterior gradient in the intraparietal sulcus (IPS) showing that the dominant factors influencing distributed numerical representations shifted from sensory modality in the posterior parietal regions to presentation format in the anterior parietal areas. Our study reveals a multifaceted brain representation of numerosity across the senses and presentation formats.

## Introduction

Whether we encounter five pens on the table, hear five knocks on the door, or hit five strikes of a hammer on a nail, we can automatically perceive ‘fiveness’. Such ability to seamlessly encode numerical information across the senses and presentation formats has led to the hypothesis of an abstract numerical code in our minds, wherein different types of numerical information are abstracted into a unified ‘fiveness’ by discarding irrelevant physical properties (Dehaene & Changeux, 1993; Wiese, 2003). The intraparietal sulcus (IPS) has often been proposed as the neural hub supporting this abstraction (Dehaene et al., 1998, 2004; Eger et al., 2003, 2009; Nieder, 2012; Piazza et al., 2004, 2006).

However, inconsistent findings across studies have made this argument controversial. Behavioral support for an abstract numerical mental code comes from studies showing no additional cost when comparing numerical quantities across spatial versus temporal formats (Barth et al., 2003) or from cross-modal adaptation effect (Arrighi et al., 2014). In contrast, other studies show format-dependent differences in the precision of numerical estimates (Tokita et al., 2013), suggesting modality-dependent representations. Consistent evidence for the existence of such abstract representations in the human brain remains elusive. While most studies suggesting the presence of an abstract numerical code in the human brain relied on invariant mean BOLD signals in parietal regions (mostly Intraparietal sulcus, or IPS) across sensory modalities and presentation formats (Dormal et al., 2010; Eger et al., 2003; Piazza et al., 2006), these results are not conclusive of abstraction since similar BOLD intensity could result from distinct neural activation within the same region. Indeed, experiments employing multivoxel pattern analysis (MVPA) to investigate whether numerical representations are supported by shared distributed neural patterns have produced inconsistent results, with some studies supporting shared representations (Damarla et al., 2016; Eger et al., 2009; Gennari et al., 2023; Nakai et al., 2023; Wilkey et al., 2020) and others failing to find evidence of abstraction (Bulthé et al., 2015; Cavdaroglu et al., 2015a; Cavdaroglu & Knops, 2019a; Czarnecka et al., 2023; Lyons et al., 2015).

These inconsistencies emerging from previous research suggest that the ‘abstract vs. non-abstract’ dichotomy may oversimplify the computational principles underlying numerical processing in the human brain. The process of abstraction may be more nuanced than generally thought. An important property of numerosity perception is that numerical quantities span multiple dimensions. It can be perceived through various sensory modalities, such as vision and audition; represented in distinct formats, including spatial (simultaneous) and temporal (sequential) arrangements; and encoded in different systems, like non-symbolic or symbolic forms. Numerical information within each format is often linked in our daily experience of the world. For example, when clapping after a show, the sequence of clapping is both seen and heard. However, co-occurrences or interactions across formats are relatively rare, as simultaneous presentations rarely have sequential counterparts across modalities. This raises the possibility that the brain may align numerical information along distinct dimensions rather than converge them into a fully unified abstract system. Results from a prior behavioral study from Arrighi and colleagues (2014) support this idea by showing cross-modal adaptation effect for temporal numerosity, which is adapting to auditory temporal numerosity effect the perception of visual temporal numerosity, and vice versa. In contrast, cross-format adaptation (e.g., adapting to temporal numerosity and testing spatial numerosity, or the reverse) have been found to be unidirectional. These findings suggest a shared representation for temporal numerosity across modalities but an asymmetry in the processing of spatial versus temporal numerosities. At the neural level, studies done in primate such as macaques and crows presented by numerosities presented either temporally or spatially found that format-independent representations emerged only after the encoding stage is finished and numerosities need to be memorized, while format-dependent representations dominated the numerosity encoding phase (Ditz & Nieder, 2020; Nieder et al., 2006; Nieder (2012)). The observation that temporal numerosity abstraction occurs during encoding, while cross-format numerosity abstraction arises only afterward, suggests different neural encoding mechanisms for temporal versus spatial numerical information. Yet in the human brain the processing of temporal and spatial numerosity, as well as their interaction, remain largely unexplored. Studies exploring whether abstracted numerical representation exist across the senses did not found evidence for an abstract multisensory representation code (Hofstetter et al. (2012); Cavdaroglu et al. (2015,2019), but those studies did not comprehensively investigate how numerical representation may align across the sense if sharing similar vs different presentation format (e.g., spatial vs temporal).

The intraparietal sulcus (IPS) plays a key role in representing numerosity. Neuroanatomical studies have shown that the IPS consists of distinct subdivisions, each with unique connectivity patterns and functional roles (Foster et al., 2022; Nelson et al., 2010; Sheng et al., 2025). Regarding numerosity, the discrete portion of the IPS seem to support specific functions (Castaldi et al., 2020; Chiou et al., 2023; Harvey et al., 2017a): the posterior parietal region is involved in presenting numerosity in space (Cavdaroglu & Knops, 2019b; Hubbard et al., 2009; Knops et al., 2009), while the anterior IPS is associated with finger counting (Andres et al., 2007). How the distinct subdivisions of the IPS contribute to the abstraction of numerical information presented in different sensory modalities or presentation formats has never been systematically tested.

In the current study, we aim to investigate numerical representation across multiple dimensions within a unified framework by using numerosity stimuli presented across modalities (vision and audition), formats (sequential and simultaneous) and notations (symbolic and non-symbolic). This approach allows us not only to investigate the effect of each dimension individually but also gain a broader perspective on the relationships between different types of representations. To minimize the intrinsic noise associated with representing large numerosities and avoid their representational overlap (e.g., Nieder & Miller, 2003; see Tsouli et al., 2021 for a review) –which may limit the ability to separate distinct numerosities using fMRI and their alignment across different formats and modalities-we purposedly decided to rely on a small numerical range (2 to 5). A multivariate pattern analytical approach (MVPA) was used to determine (1) whether and where the brain encodes numerical information across the senses and presentation format or notation, (2) whether those regions implement an invariant neural representation across different dimensions and sensory input and (3) elucidate the neural organization of numerical representation across dimensions using representational similarity analyses (RSA) combined with computational models.

## Results

Participants engaged in a numerosity one-back task in which four numerosities from five different stimulus conditions span (see below) were presented in a pseudo-randomize order. The sequence was designed so that the same stimulus condition could not be presented in consecutive trials. Participants were asked to compare the current numerosity with the previous one, regardless of the condition formats or modalities, and to press a button with their right index finger when the stimulus in the current trial had the same numerosity as the previous trial (target stimuli, 10% of the trials; Figure 1A). Participants were instructed to avoid active counting during the task.

**Figure 1.**
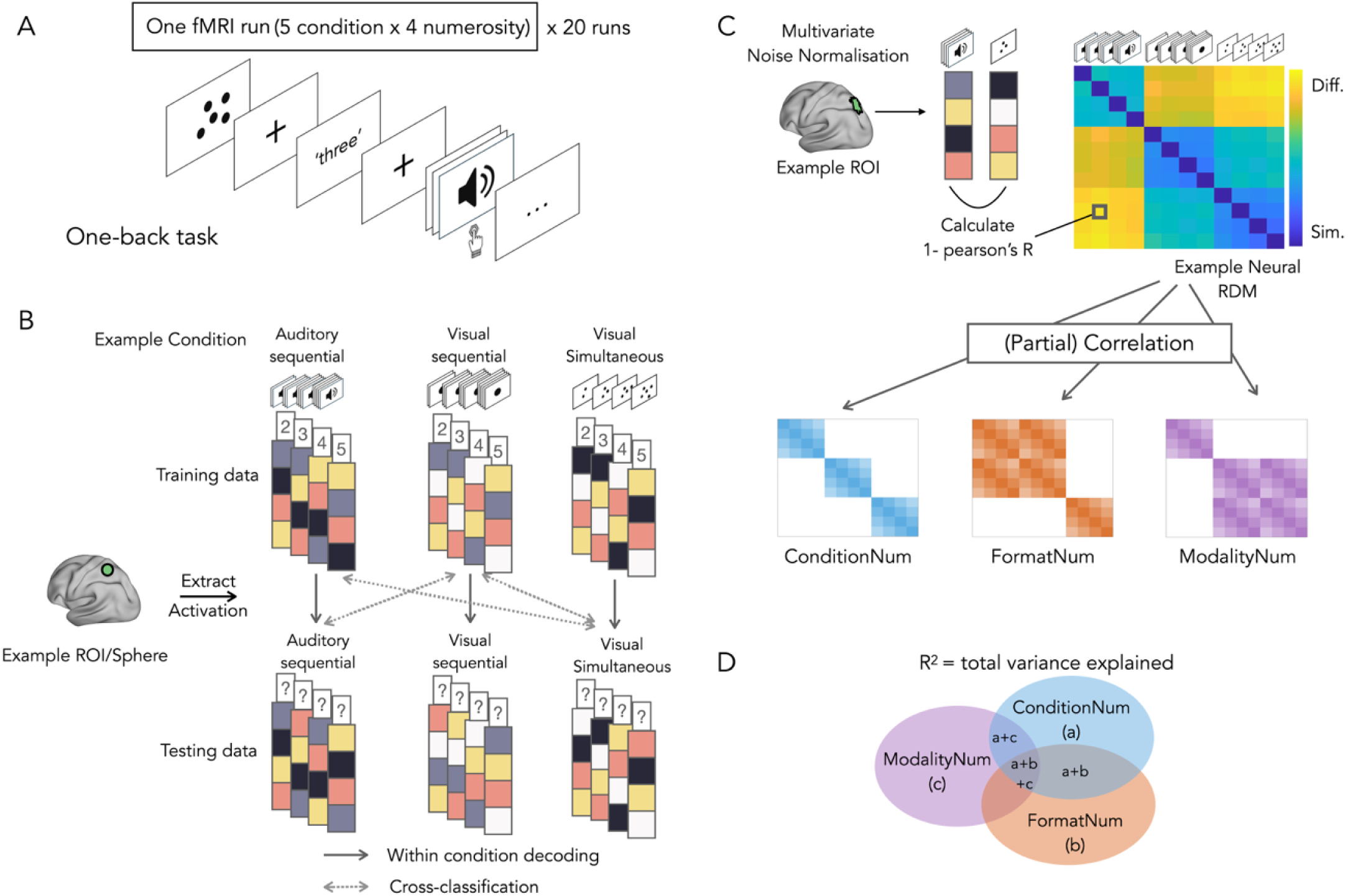
Experimental design and data analysis scheme. **A.** experiment task. During the fMRI session, Participants were instructed to press a button whenever the quantity of the current numerosity matched that of the previous one, regardless of format or modality. **B.** For decoding analysis, voxel-wise neural activation patterns were extracted for each region of interest (ROI) in the ROI-based analysis, or for each searchlight sphere in the searchlight analysis. Pattern vectors were divided into training and testing sets. For within-condition decoding, both sets were taken from the same condition. For cross-classification, training and testing sets were from different conditions. **C.** For RSA analysis, multivariate noise normalization was first applied to the beta coefficients obtained for each stimulus. Then for each ROI, a neural representational dissimilarity matrix (RDM) was constructed by computing pairwise dissimilarities between condition vectors (1 – Pearson correlation). Three theoretical model RDMs were generated based on specific hypotheses. Correlations and partial correlations were computed between each model RDM and the neural RDM. **D.** The Euler diagram illustrates the results of the variance partitioning analysis. The total explained variance of the full model is indicated by R^2^. Each region of the diagram represents the proportion of variance uniquely or jointly explained by the individual model components.

Numerosities 2, 3, 4, 5 using visual sequential, auditory sequential, visual simultaneous, visual symbolic (Arabic), auditory symbolic (verbal number words) conditions. Visual simultaneous and visual sequential numerosities were presented on a black background using white dots. Visual symbolic numerosities were presented as white Arabic numerals on a black background. The non-numerical sensory features of non-symbolic stimuli were balanced using two types of stimuli (see Methods for details). The behavioral results showed overall good performance (localizer task: mean accuracy = 83%, ranging from 68% to 91%; decoding task: d-prime score = 3.2102 on average, ranging from 1.0683 to 4.9301), suggesting that participants were actively attending to the experiment.

## fMRI results

### Whole brain searchlight decoding analysis

To get an overview of brain regions involved in representing numerosities across conditions, we conducted a whole-brain searchlight analysis for each condition. The resulting maps corresponding to all stimulus conditions are shown in Figure 2. The auditory sequential and visual sequential conditions exhibited similar patterns, with significant decoding observed across a widespread network that includes the frontal, parietal, occipital and temporal areas. For the visual simultaneous condition, significant decoding was observed in the occipital, parietal and frontal cortices. In contrast, the two symbolic conditions exhibited much less, if any, significant decoding. In the auditory symbolic condition, numerosity decoding could only be found in small cluster of right superior temporal cortex. No significant voxels were identified in the searchlight results for the visual symbolic condition, and thus, no results map was presented for this condition. Therefore, due to the limited or absent decoding observed in the symbolic conditions, we excluded them from further analysis.

**Figure 2.**
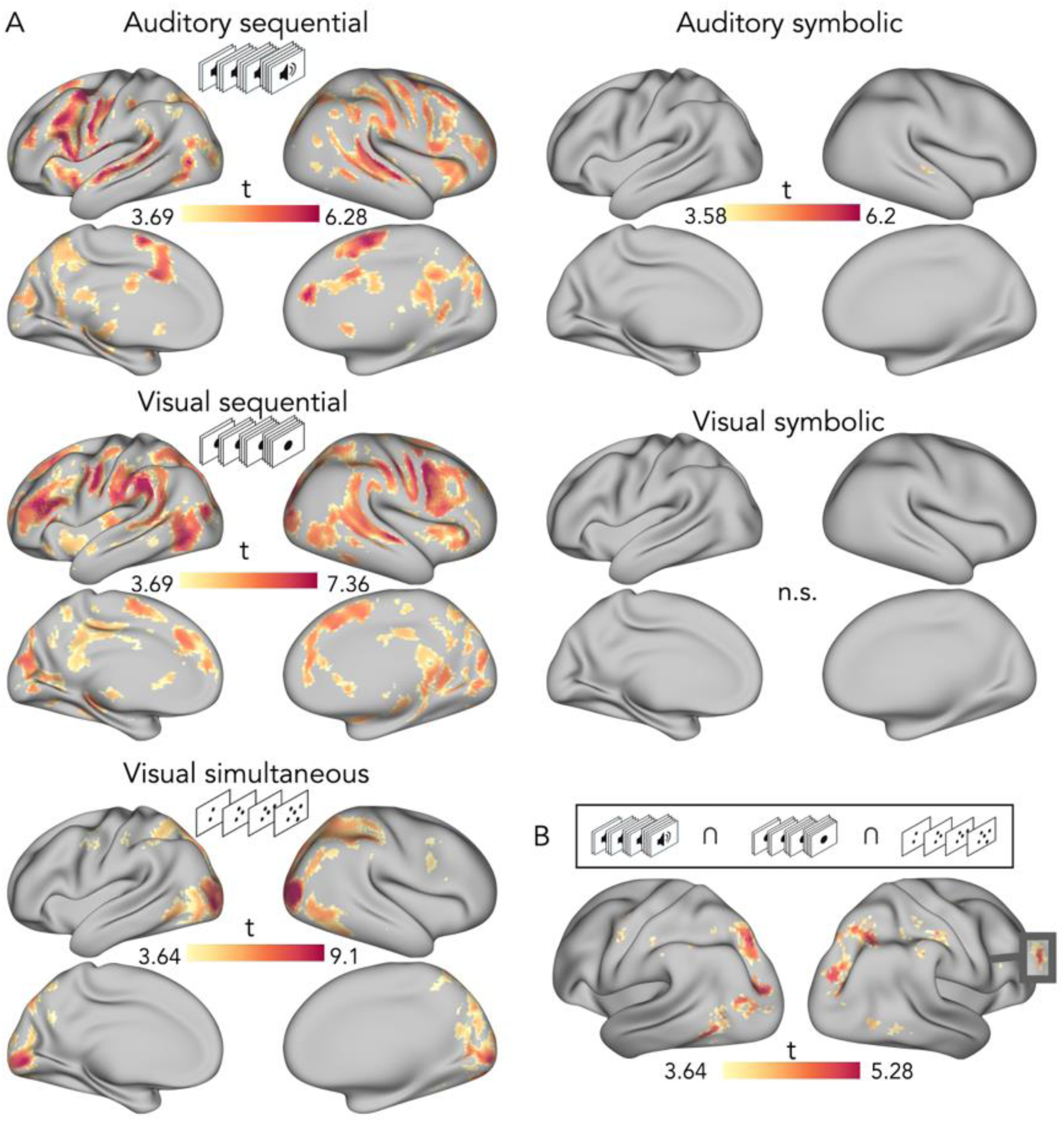
**A**: With-in condition Searchlight numerosity decoding from different conditions. P values were set at p < 0.001 for cluster-level correction with probabilistic threshold-free cluster enhancement (pTFCE). **B**: The results map for the conjunctions of the searchlight maps for three non-symbolic conditions (i.e., auditory sequential, visual sequential, visual simultaneous conditions) with a threshold of p< 0.05 FEW corrected at cluster level.

To localize brain regions consistently engaged in decoding numerosity across all non-symbolic conditions, we conducted conjunction analyses (Nichols et al., 2005) of the searchlight maps for the auditory sequential, visual sequential and visual simultaneous conditions. The results revealed overlapping regions along the dorsal stream, extending from lower-level occipital areas to the parietal and frontal regions, which were engaged in decoding numerosity across all non-symbolic conditions (Figure 2B). Among these regions, a total number of seven clusters showed significant decoding with FEW-correction (.05) at cluster level and were thus identified as regions of interest (ROIs) for further analysis (tableS3). One ROI was in the visual cortex, five in the parietal cortex, and one in the frontal cortex (Figure 4A). To ensure that successful decoding was not driven by low-level features specific to one subset of stimuli, we conducted a control analysis in which a classifier was trained on data from both subsets but tested only on one subset. The control analysis showed above-chance level decoding accuracy when a classifier was tested on data from only one subset for all three non-symbolic cinditions. This suggests that the successful decoding was not fully driven by low-level features. (Figure S4).

## ROI-Based Analyses

### Distance effect

ROIs were grouped into three cortical regions: occipital, parietal and frontal ROI regions and the numerical distance effect, a hallmark of magnitude encoding, was examined for each condition in each group. The numerical distance effect was observed across all conditions in the parietal region and in most conditions in the frontal region (Figure 3). Specifically, in the parietal region, decoding accuracy increased linearly with numerical distance across all conditions (auditory sequential: t(61) = 3.90, p_corr_ < 0.001, Cohens’*d* = 0.60; visual sequential: t(61) = 4.47, p_corr_ < 0.001, Cohens’*d* = 0.60; visual simultaneous: t(61) = 4.06, p_corr_ < 0.001, Cohens’*d* = 0.62). In frontal region, significant linear increases in decoding accuracy were observed for the auditory sequential and visual simultaneous conditions (auditory sequential: t(61) = 2.72, p_corr_ = 0.025, Cohens’*d* = 0.42; visual simultaneous: t(61) = 3.42, p_corr_ = 0.003, Cohens’*d* = 0.53), but not for the visual sequential condition. No significant linear increase in decoding accuracy was observed in the visual region.

**Figure 3.**
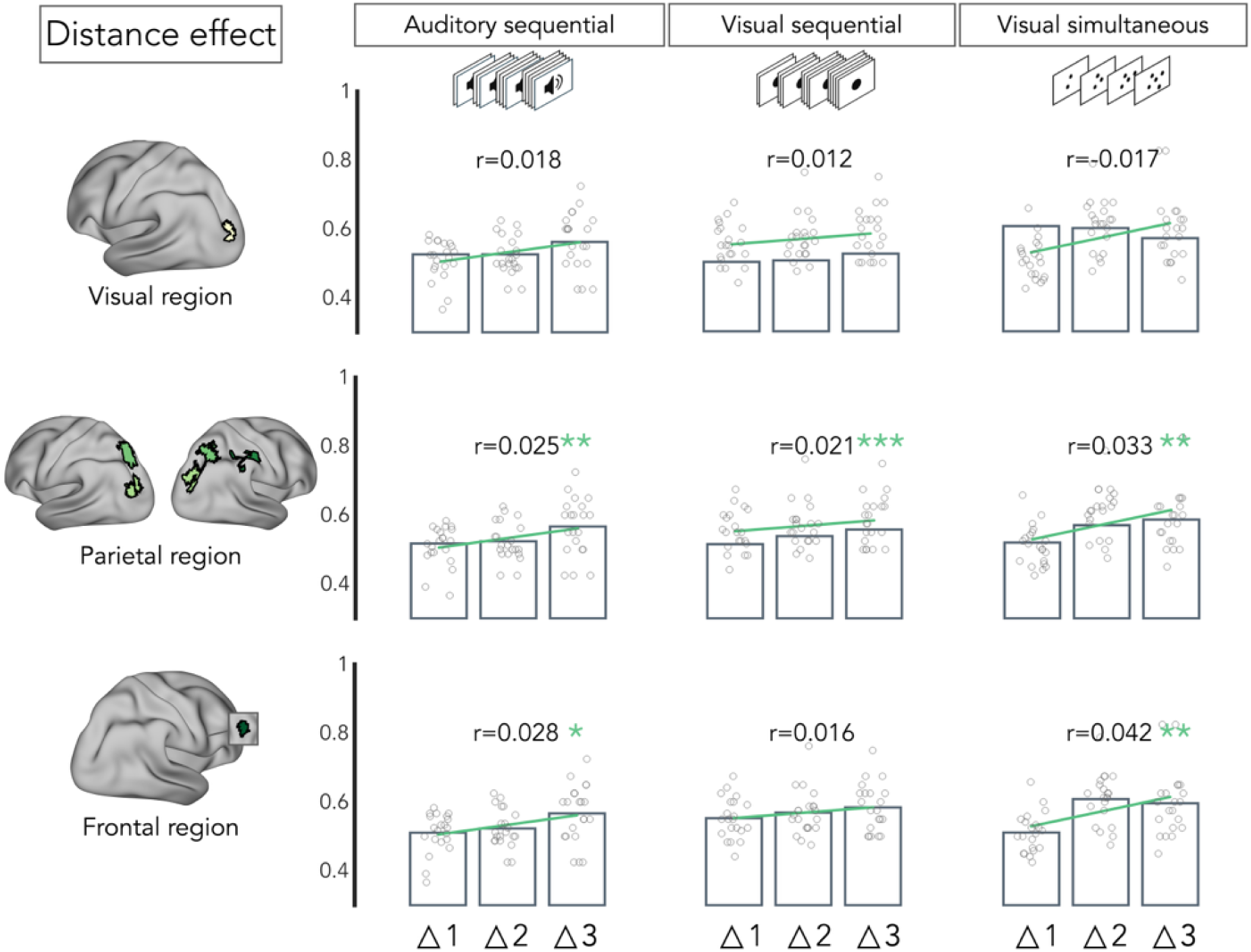
Distance effect for ROIs in each cortical region. Results are FDR corrected for three conditions for each region (*p < 0.05, **p < 0.01, ***p < 0.001).

### Cross-classification Analysis

To identity whether ROIs that can decode numerosity across all non-symbolic conditions (i.e., ROIs identified through conjunction analysis) exhibit invariant representation patterns for numerosities, we conducted cross-classification analyses where the classifier was trained on numerosities from one condition and tested on numerosities from another condition and reversely (e.g., training on visual sequential condition and testing on the auditory sequential condition, and vice versa; FDR-corrected). The decoding accuracy reported here represents the average across both directions of training and testing. For visual sequential versus auditory sequential pairs, significant cross-classification results were observed in all the parietal ROIs (LhIP4, p = 0.0124, LhIP8, p = 0.0104, RhIP2, p < 0.001, RhIP3, p = 0.0019, RhIP4, p = 0.0099) and frontal ROIs (Precentral, p < 0.001) but not in the visual cortex (LV3A, p = 0.2423) (Figure 4B). Decoding accuracy for each individual direction is provided in the supplementary material (Figure S2).

**Figure 4.**
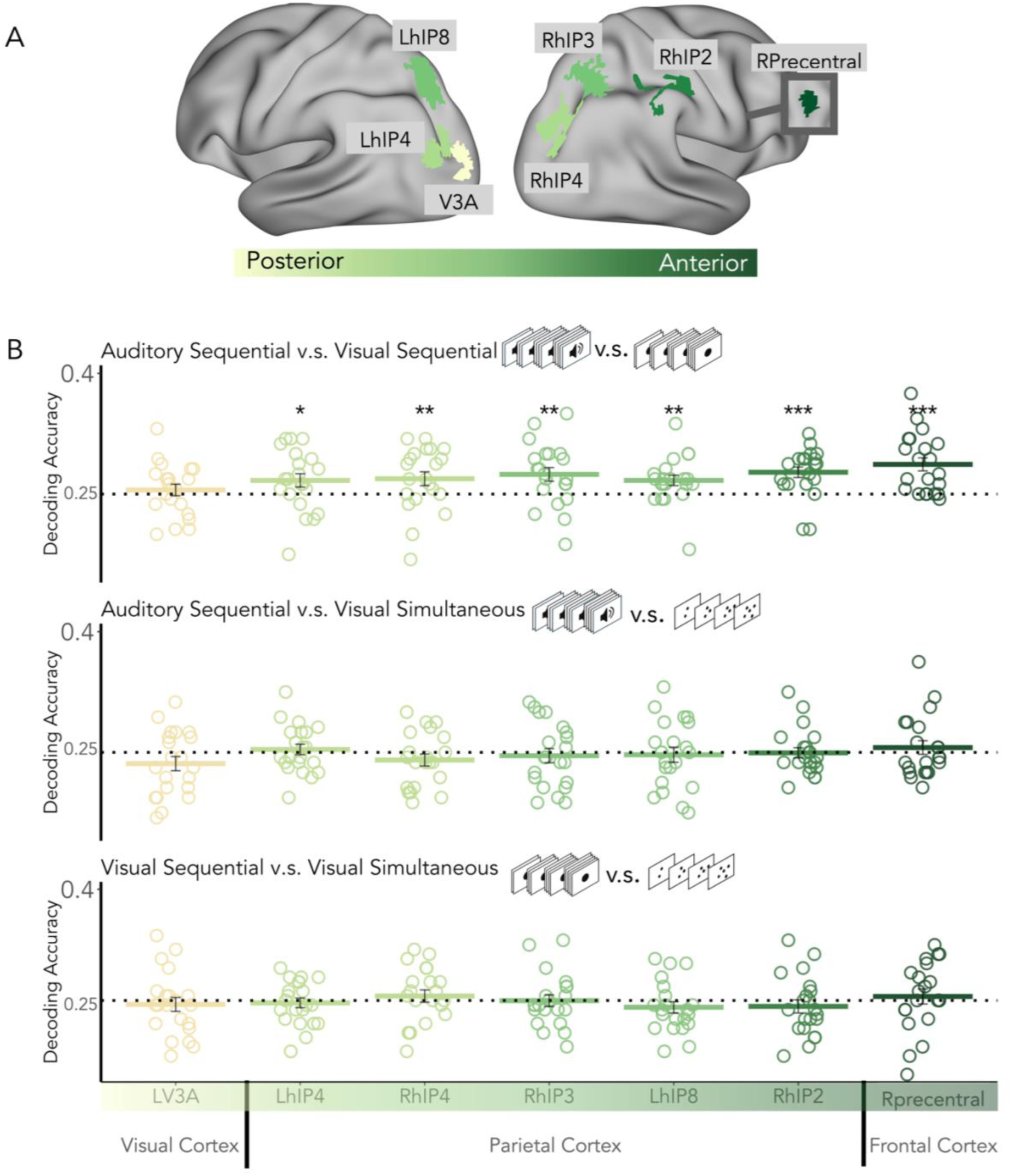
**A**: ROI that were identified from conjunction map. ROIs are illustrated as colored patches, with color progressing from light green to dark green along the cortical surface from posterior to anterior. **B**: Cross-classification results in ROIs illustrated in Figure 3B. Chance level (dotted line) is 25%. Individual points reflect individual subjects’ results. Results are FDR corrected (*p < 0.05, **p < 0.01, ***p < 0.001). Error bars represent standard error of mean.

For both auditory sequential versus visual simultaneous pairs and visual sequential versus visual simultaneous pairs, no significant cross-classification was observed across any of ROIs, either in the average of the two directions or in each direction separately (Figure 4B, Figure S2).

### Representational similarity analysis

RSA analyses complement cross-classification decoding by moving beyond a simple dichotomy of shared versus no-shared information to a more comprehensive exploration of the geometry of numerical representation. By using RSA and theoretical models, we were able to further quantify the similarities and differences in numerical representations across conditions and modalities.

First, we observed a numerically high and similar upper-bound level of the noise ceiling across fROIs regions showing high inter-subjects consistency in how these regions react to our stimuli space. Interestingly, some models were close to this noise ceiling level according to the region tested, supporting their validity.

We built three theoretical models that we thought could represent how brain regions encode our different numerosities, the Condition-Numerosity (CondNum) model assumes greater similarity of numerosities within conditions, the Modality-Numerosity (ModalNum) model assumes greater similarity within the same modality across formats, and the Format-Numerosity (FormatNum) model assumes greater similarity within the same format across modalities (see Figure 1C). We then tested the correlation between the neural dissimilarity matrix (DSM) of each ROI in each subject with three theoretical models (Figure 5 A, B). All three models are significantly correlated with neural DSM across all ROIs [all: W(21) ≤ 21, r_bc_ ≤ 0.0909, p ≤ 0.001].

**Figure 5.**
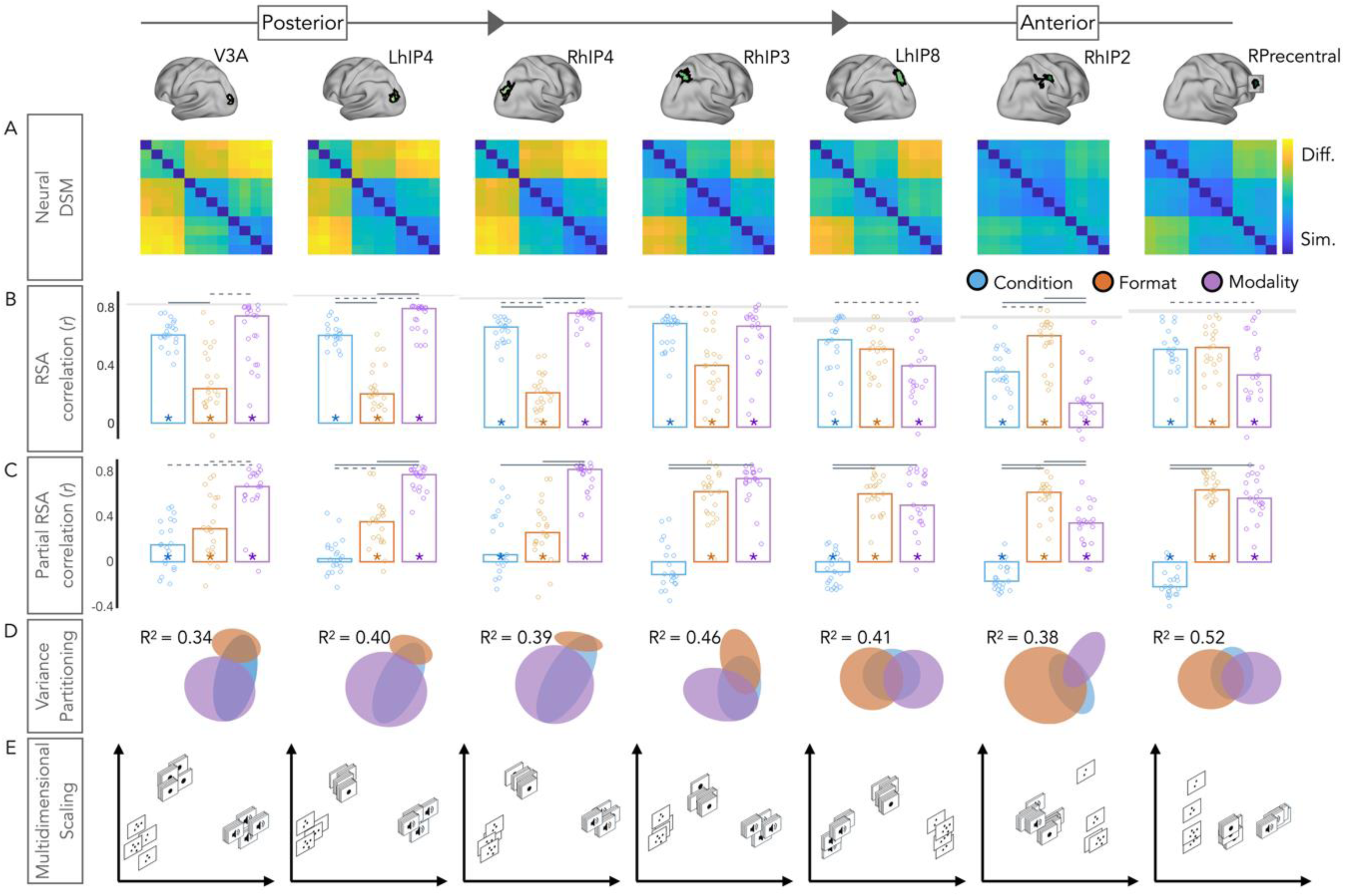
**A**: Group-averaged neural RDMs. **B**: Bar charts indicating the median Spearman’s r rank-order correlations between neural RDMs for individual subjects (dots) and model RDMs. Gray blocks indicate the noise ceiling; Stars mark models whose correlations differ significantly from zero (one-sided Wilcoxon signed-rank test, p < 0.05). Connecting bars show two-sided pairwise comparisons between model correlations: solid bars indicate p < 0.001, dashed bars indicate p < 0.05. **C**: partial correlation for each model RDM. Stars mark models whose correlations differ significantly from zero (one-sided Wilcoxon signed-rank test, p < 0.05). Connecting bars show two-sided pairwise comparisons between model correlations: solid bars indicate p < 0.001, dashed bars indicate p < 0.05. **D**: Euler diagram depicting results of variance partitioning the individual dissimilarity in each ROI between models. **E**: Two-dimensional multidimensional scaling applied to the dissimilarity matrices. Stimulus modality and presentation format are indicated by shape: circles represent auditory sequential numerical stimuli, crosses represent visual sequential stimuli, and triangles represent visual simultaneous stimuli. The size of each shape scales with numerosity

However, the relative contributions of the models varied across ROIs. A gradual shift was observed from posterior to anterior areas: the contribution of the modality numerosity model decreased while that of the format numerosity model increased. More precisely, in the early visual cortex and several posterior parietal subregions, the modality numerosity model exhibited the strongest correlation with neural patterns while in RhIP2, the most anterior parietal subregion examined, the format numerosity model showed the highest correlation. Detailed statistical results are provided in the Supplementary Material.

### Partial correlation and variance partitioning

To explore independent contribution of each model to the neural activity, we computed partial correlations between the models and neural activities (Figure 5C). In general, although the condition numerosity model was significant in most fROIs, its contribution was consistently lower than that of the modality and the format models, suggesting that these two features of stimuli presentation were crucial in representing how numbers are encoded in these regions. The modality numerosity model and format numerosity model were both significantly correlated with neural DSM across all ROIs [all: W(21) ≤ 31, r_bc_ ≤ 0.1342, p ≤ 0.002].

Importantly, we observed a dissociation across ROIs regarding the relative contributions of the modality and format models. A gradual shift was observed from posterior to anterior areas: the contribution of the modality numerosity model decreased while that of the format numerosity model increased. Specifically, in the early visual cortex V3A and posterior parietal subregions left hIP4 and right hIP4, the modality model contributed the most to the neural activity, with significantly stronger effects than the format model [all: W(21) ≤ 46, r_bc_ ≤ 0.1991, p ≤ 0.005]. In the lateral parietal subregions left hIP8 and right hIP3, the contributions of the modality and format models were not significantly different [left hIP8: W(21) = 115, r_bc_ = 0.4978, p = 0.2305; right hIP3: W(21) = 62, r_bc_ = 0.2684, p = 0.0537]. In contrast, in the anterior parietal subregion left hIP2, the format model contributed the most, showing a significantly greater influence than the modality model [W(21) = 25, r_bc_ = 0.1082, p ≤ 0.001]. Finally, in the frontal Precentral region, the contribution of modality model and format models did not differ significantly [left hIP8: W(21) = 99, r_bc_ = 0.4286, p = 0.0735]. Detailed statistical results are provided in the Supplementary Material.

When all three predictors were regressed on the neural dissimilarity matrix, the full model explained significant portion of the variance in the group neural RDMs (all: R^2^ > 0.338, p<0.001). However, the distribution of explained variance varied across ROIs (Figure 5D). In the early visual cortex and posterior parietal subregions (LhIP4 and RhIP4), the modality model uniquely explained large portions of the variance. The MDS plot reveals a clear separation of modality across conditions (Figure 5E). In the lateral parietal subregions (LhIP8 and RhIP3), both modality and format models contribute substantively to the explained variance. In contrast to the posterior parietal subregions, the format model was the primary contributor to explain variance in the anterior parietal subregions. The MDS plot show a clear division between format across conditions. In frontal regions, the format models remained the stronger contributor, though its advantage over the modality model was less pronounced compared to the anterior parietal subregions. Overall, these variance partitioning results corroborate those observed with RSA.

## Discussion

Humans and other animals seamlessly extract numerosity from a countless variety of situation. Whether the human mind and brain implements an abstract numerical code has been debated for the last two decades (Cohen Kadosh & Walsh, 2009). In this study, we examined how numerical representations are organized across different sensory modalities and representation formats. Relying on multivariate decoding we first aimed to identify regions that encodes numerical information in their distributed activity patterns across the senses and presentation format. We first reveal a set of overlapping brain regions, spanning from V3A to multiple IPS subregions and precentral area in prefrontal cortex that successfully encode numerical information in all three non-symbolic conditions. These results confirm the crucial role of a dorsal occipito-parieto-frontal network in numerical processing (Castaldi et al., 2019; Dehaene et al., 2003; Harvey et al., 2017b; Knops, 2017; Kido et al., 2025), extending its role to reliable encoding of numerosities across senses and formats. Cross-classification analysis further revealed a shared numerosity code across vision and audition when numerosities are presented using a similar temporal code. Finally, using representational similarity analyses, we identified a posterior-to-anterior gradient in the IPS with numbers being encoded in posterior regions primarily relying on the senses while anterior region coding more for the format of numerosity presentation.

Numerosity is inherently embedded within both spatial and temporal domains, each engaging distinct cognitive mechanism. While simultaneous numerosity relies on visuospatial processing and are parallelly extracted through location maps (Anobile et al., 2018; Dehaene & Changeux, 1993), sequential numerosity unfolds over time, requiring serial accumulation and engaging temporal processing. Previous studies have suggested that parietal and frontal cortices perform a ‘normalization’ process in which numerosity representations are abstracted into a unified representation by discarding irrelevant physical properties (Croteau et al., 2024; Dehaene & Changeux, 1993; Park & Huber, 2022). However, our findings suggests that this normalization process does not entirely eliminate physical information but instead maintains a distinction between spatial and temporal numerosity representations. Indeed, our cross-classification analysis showed that in parietal and frontal cortex, there is an invariant neural pattern for numerosity representation across visual and auditory modalities but only when sharing a sequential representation format. This shared representation does not extend across different presentation formats: neural patterns for numerosity in sequential (visual/auditory) formats do not align with those from simultaneous (visual) presentations. In other words, even in regions that both code for spatial and temporal numerical information, no alignment is found across those two formats of numerical displays, even if presented within the same modality (here, vision). This result can be understood by daily statistics since numerosities within each presentation format – and especially the sequential format – tend to be closely linked to each other and often co-occur. In contrast, co-occurrence or interactions across different presentation formats is rare. The brain may internalize these statistical patterns from everyday experience (Qu et al., 2024; Zhao & Yu, 2016), classifying numerosities from the same presentation formats more closely to one another even if coming from different senses. From an efficiency perspective, generalizing numerical representations within the temporal domain while preserving a distinction between spatial and temporal representations may serve to optimize processing efficiency and reduce inter-domain interference (Czajko et al., 2024).

We carefully controlled low-level features for numerosities from each condition by using two sets of stimuli. We further restrict our cross-classification analysis on brain regions that successfully decode all non-symbolic numerosities to exclude brain regions sensitive to specific low-level features. However, since the cross-classification only succeeds when numerosity shared a similar temporal code, one may still question whether this cross-classification could be driven by low-level temporal features. This is unlikely for several reasons. First, total sequence duration, individual beep/dot duration and inter-beep/dot intervals were varied across the two stimulus subsets, making it difficult to rely on any of these features to discriminate numerosity (Cavdaroglu et al., 2015b; Dehaene et al., 2005). Second, both the individual beep/dot durations and inter-beep/dot intervals were jittered to prevent any regular patterns and participants relying on the frequency of stimuli as a cue to discriminate numerosity. A prior study where participants maintained a 6-second tactile vibration or visual flicker with embedded frequency information did not identify a distributed supramodal frequency representation in parietal regions (Dakin et al., 2011). Our cross-classification results in a set of parietal subregions suggest that it may be the extraction of numerosities, rather than frequency, that facilitates the alignment of sequentially presented numerosities across the senses.

Our findings align with a previous study showing that number-selective neurons in the prefrontal and parietal cortices of non-human primate encodes sequential numerosity across sensory modalities (Nieder, 2012). To our knowledge, the current study is the first to reveal such an alignment in the human brain. While Cavdaroglu et al., (2015a) previously attempted to examine cross-modal alignment of sequential numerosity in the human brain using similar cross-classification approach, their results did not support the existence of an alignment. In contrast, our results provide unequivocal evidence of cross-modal alignment for sequential numerosity. Compared to Cavdaroglu et al.’s study, which used a broad range of numerical stimuli ranging from 5 to 16, we intentionally relied on a small number range in order to trigger a salient representation of specific numerosities, something unlikely with larger numbers since their internal representations become noisier, less precise and less accurate (Tokita et al., 2013). This in turn may reduce the brain’s capacity to create a unified representation of larger numerosities across different stimulus types or modalities. Furthermore, our study shows that an active response is unnecessary for detecting numerosity representations or their alignment (Cavdaroglu et al., (2015a)). Our experiment relied on a one-back task requiring participants to memorize previous quantity and discriminate if the ongoing quantity is the same as the previous one. Response was not needed for most of the trials. One may argue that the activation of a verbal code during the comparison phase might be necessary for aligning numerosity representations. However, our results argue against this interpretation since alignment occurred only when numerosity shared a representation format. Furthermore, the structure of the task was identical for symbolic numerosities for which we did not observe reliable decoding. Altogether these results suggest that the activation of a verbal code does not underly our observation of a shared temporal numeral code across the senses, but rather the presence of a shared representation format that ultimately determines alignment.

In the current study, we did not find any invariant neural patterns across presentation formats, that is, across the sequential and simultaneous presentation formats. Nieder’s study examined number-selective neurons for visual numerosity across these formats in the non-human primate brain and found that when the enumeration process was completed and animals needed to maintain the quantity in mind for an upcoming quantity-matching task, a subset of neurons (19 % of recorded neurons) encoded the quantity irrespective of the presentation format (simultaneous, sequential). Although our participants also need to retain the quantity in memory, we did not observe an invariant neural activity pattern across presentation formats. One potential explanation for this discrepancy is that only a small group of neurons exhibit such shared representation, potentially impeding the ability of classic fMRI to detect such restricted signal in space. Alternatively, the limited temporal resolution of fMRI makes it challenging to capture fast and transient changes in neural activity in a timely manner as was done by Nieder and colleagues’ study who could see that across format alignment does not express during the encoding but only during the maintaining phase (Ditz & Nieder, 2020; Nieder et al., 2006). Such abstraction process might thus require a finer level of spatial and temporal resolution to be successfully assessed, not easily accessible with classic fMRI. Future research using ultra high-field fMRI combined with techniques having temporal sensitivity (e.g., i/M/EEG) could provide deeper insights into the presence of an abstract representation across presentation formats in the human brain.

A notable finding in the present study is the successful decoding of non-symbolic numerosity, including auditory sequential numerosity, in V3A. The extent to which early visual cortex supports numerosity processing remains debated. People supporting this idea argue that numerosity is a primary perceptual property, which can be perceived directly (Castaldi et al., 2019, 2021; Cicchini et al., 2016; DeWind et al., 2019; Fornaciai & Park, 2018; Park et al., 2016; Testolin et al., 2020), while others suggest that numerosity can only be perceived indirectly through other visual cues (Dakin et al., 2011; Durgin, 2008; Gebuis et al., 2014; Paul et al., 2022). Our results contribute to this debate by showing that, beginning in V3A, the early visual cortex is able to extract numerical information from a variety of sensory inputs (Fornaciai & Park, 2018). Moreover, the successful decoding of auditory sequential numerosity in V3A indicates that its function extends beyond the processing of spatial visual features. However, the numerosity representation in V3A appear to remain tied to their sensory origin rather than being abstracted.

Surprisingly, we did not observe any reliable decoding for visual or auditory symbolic numerosity across the whole brain. The very strong decoding in other conditions refute the idea this null result relates to the sensitivity of our paradigm, fMRI acquisition parameters or analytical scheme. A prior study compared visual symbolic numerosity, dot arrays and braille numerosities and neither did find representations of visual symbolic numerosity (Czarnecka et al., 2023). They hypothesized that symbolic numerosity, due to overtraining, evoked lower activation and therefore lower “decodability”. In the current study, univariate analyses for the symbolic conditions evoked comparable univariate activation level when compared to the non-symbolic conditions ruling out such interpretation (see supp. Figure S1). Another possibility would be that symbolic numerosity representations may require higher spatio-temporal sensitivity to be detected. This idea is consistent with the hypothesis that non-symbolic numerosity, as the evolutionarily older one, is probably represented by a larger neuronal population (Ansari, 2008; Eger et al., 2009) that is potentially innate (Lorenzi et al., 2025).

By constructing theoretical models and correlating them with distributed neural activity patterns using RSA, we further unravelled a posterior-to-anterior organization gradient along the dorsal pathway. In the most posterior part of the IPS and V3A, numerical representations are mainly encoded in terms of sensory inputs while the most anterior IPS is primarily encoding numbers in terms of their presentation format. Intermediate regions, including the lateral IPS, exhibit a comparable influence of both model and may function as a transition hub from modality to format processing. Unlike the cross-classification results, which served as a more stringent approach to address the presence of partially abstracted numerical representation in a region, RSA provides a finer-grained measurement of a progressive transition from primarily modality-based to primarily format-based processing along the parietal cortex. The posterior IPS regions are highly connected with extrastriate visual areas (Bray et al., 2013). In contrast, hIP2, the most anterior IPS subdivision, situated next to premotor cortex, exhibits a close link with the ventral premotor cortex (Uddin et al., 2010). This differential connectivity profile suggests that visual sensory inputs processed in posterior IPS regions are transformed into motor actions through connections from the anterior IPS to the prefrontal cortex. Indeed, studies in non-human primates have found that posterior IPS had the strongest functional connectivity to regions involved in motion processing and eye movements, while anterior IPS was functionally connected to regions associated with motor, tactile, and proprioceptive processing, as well as regions involved in reaching, grasping, and processing peripersonal space (Knops et al., 2009; Sheng et al., 2025). In numerosity studies, the anterior IPS also show a linkage between numerical perception and motor functions, exemplified by its role in encoding finger counting (Kaufmann et al., 2008; Krinzinger et al., 2011). We therefore speculate that the temporally structured organization in the anterior IPS region hIP2, given its anterior position and link to the premotor cortex, may serve for supporting motor actions (Anobile et al., 2016; Kirschhock & Nieder, 2022; Sawamura et al., 2020; Togoli et al., 2020). Recently, researchers have proposed a sensorimotor theory of numerosity processing, indicating a more generalized system in the brain that integrates action with the processing of other quantitative magnitudes such as space and time (Anobile et al., 2021; Sixtus et al., 2023). Our results may reflect this sensorimotor organization of numerosity, suggesting that the brain automatically aligns numerosity information based on its temporal structure. Such an organization may facilitate motor actions, as action unfold over time, making their numbers embedded in their sequential nature. The anterior IPS region hIP2, where we found stronger format-based coding, may therefore serve as a key locus in this sensorimotor organization.

In conclusion, our study investigated how numerical representations from various sensory modalities and formats converge into a higher-level numerical representation in the brain, revealing a multifaceted organization of numerosity across modalities and formats. We observed a posterior-to-anterior gradient organization along the dorsal pathway, beginning with numbers encoded mostly based on their sensory attributes in early visual cortex, progressing with a mixture of processing of sensory and format information in the posterior parietal cortex and ending to a primarily format-based processing in anterior parietal cortex. This distributed gradient encoding offers critical insights into the brain’s mechanisms dedicated to numerical processing from various sensory inputs and format of presentation. Our study suggests that numerosity might not converge into a single, generalized representation abstracted from all physical properties. Instead, it appears to preserve a distinction between spatial and temporal numerosity representations, possibly reflecting statistical regularities in daily experience and putatively contributing to more efficient neural processing. This temporal organization of numerosity becomes most salient in anterior parietal, maybe playing a role in transferring numerosities into a corresponding action effectively.

## Methods

### Participants

A total of twenty-three individuals participated in the experiment, completing two 90 minutes fMRI scanning sessions on two different days. The mean age of the participants was 23.69 years (SD = 3.46), with an age range of 19 to 35 years. All participants were right-handed and had normal or corrected-to-normal vision. Participants reported no history of neurological conditions. Two subjects were excluded from data analysis due to either lack of response during fMRI acquisition, suggesting no attention to the task, or incorrect acquisition parameters. The protocol was approved by the research ethics committee in Saint-Luc University Hospital (protocol number 45.327.621). Written consent was acquired from all participants before participation. Participants were monetarily compensated for their participation. The sample size of the experiment was determined a priori based on similar fMRI MVPA study on numerosity representations (Cavdaroglu et al., 2015a; Cavdaroglu & Knops, 2019a; Czarnecka et al., 2023).

### Numerosity decoding experiment and stimuli

Participants engaged in a numerosity one-back task in which four numerosities from five different stimulus conditions (see below) were presented in a pseudo-randomize order. The sequence was designed so that the same stimulus condition could not be presented in consecutive trials. Participants were asked to compare the current numerosity with the previous one, regardless of the condition formats or modalities, and to press a button with their right index finger when the stimulus in the current trial had the same numerosity as the previous trial (target stimuli, 10% of the trials). Participants were instructed to avoid active counting during the task.

Numerosities 2, 3, 4, 5 using visual sequential, auditory sequential, visual simultaneous, visual symbolic (Arabic), auditory symbolic (verbal number words) conditions. Visual simultaneous and visual sequential numerosities were presented on a black background using white dots. Visual symbolic numerosities were presented as white Arabic numerals on a black background.

To minimise the possibility that participants encode numerosity based on low-level stimuli features, the non-numerical sensory features of non-symbolic stimuli were balanced using two types of stimuli. For visual simultaneous stimuli (arrays of dots), intensive parameters (individual item size and density) are controlled for one set while extensive parameters (total area and convex hull) are controlled for the other set (Dehaene et al., 2005). Stimuli were generated using GeNEsIS, a software platform designed for creating visual numerical arrays with precise control over continuous variables (Zanon et al., 2022). The average individual dot size, convex hull, total area and density for each set and numerosity can be found in supplementary materials (Table S1). Stimuli were displayed on an area of approximately 4.8 × 4.8 visual degree. The low-level features of auditory sequential stimuli (sequential beeps) were similarly controlled by 2 sets. For one set the total sequence duration is fixed while for the other set the individual beep duration is fixed. Individual beep durations and inter-beep intervals were jittered to avoid any regular patterns that might predict numerosity and interfere with any counting strategy (Kanjlia et al., 2021). The audio frequency of each individual beep was 440 Hz. The auditory stimuli were prepared using custom MATLAB scripts (R2019b; MathWorks). The duration of individual beeps was designed to be less than 270 ms to prevent counting (Cavdaroglu et al., 2015a; Piazza et al., 2006; Tokita & Ishiguchi, 2011). However, for the set that the total sequence duration is fixed, it is impossible to keep individual beeps shorter than 270 ms especially for small numerals such as 2 and 3. Therefore we use beeps longer than 270ms for them. The single beep durations and total stimulus duration are provided in supplementary materials (Table S2). The visual sequential stimuli were generated based on the same logic applied to auditory sequential stimuli. However, during piloting, we observed that visual sequential stimuli were more challenging to discriminate compared to auditory ones. To ensure comparable difficulty levels across conditions, we increased the total duration to 1.2 times longer than auditory stimuli while maintaining the internal structure unchanged. Visual sequential stimuli are presented on a black background using white dots, spanning about 0.73 visual degrees. The single dot durations and total stimulus duration are provided in supplementary materials as well (Table S2). In visual symbolic conditions, the position and size of digits on the screen varied across trials to prevent adaptation to low-level features, with the position varying within a range of 1.21 visual degrees. In auditory symbolic conditions, number words for each numerosity were presented using two different voices—one male and one female—to avoid adaptation effects.

The experiment comprised 20 runs, containing 40 trials (5 conditions × 4 numerosity × 2 sets of condition controlled for low-level features) and four additional target trials (44 trials in each run). The inter-stimulus interval between each stimulus ranged from 3.5s to 4.5s. Auditory stimulation was delivered through in-ear MRI-compatible earphones (Sensimetric S14, SensiMetrics, USA) at a loud but comfortable volume, defined with each participant before the start of the MRI acquisition. Visual stimuli were presented on a screen through a mirror at a distance of 170 cm and with a resolution of 1920×1200 pixels. To ensure fixation and to minimize eye movements, a fixation cross was shown in the center of the screen throughout each run, except during the presentation of the visual sequential stimuli, where flashing dots were presented at the center of the screen. Visual simultaneous stimuli were presented for 1.25s. Auditory sequential stimuli had a total duration between 0.55s and 1.6s. Visual sequential stimuli had a total duration between 0.59 to 1.74s. Visual symbolic stimuli were presented for 1s, and auditory symbolic stimuli were played for 500ms. The stimulus delivery was controlled using the Psychophysics toolbox implemented in Matlab R2019b (The MathWorks).

## Behavioral Data Analysis

In order to characterize the behavioral performance, we used the *d’* measure (Snodgrass & Corwin, 1988) instead of the simple accuracy rate. This is because accuracy rate only accounts for hit or miss target trials (10%), making it one-sided. Instead, the *d’* not only considers the hit rate of target trials but also penalizing for responses following non-target trials (false alarm). The *d ′* is calculated from hit rate and false-alarm (FA) rate using the formula *d ′* = Z_Hit_ – Z_FA_.

## fMRI data Acquisition and preprocessing

Functional images were acquired at the Saint-Luc University Hospital, Brussels, Belgium with 3T GE scanner (Signa™ Premier, General Electric Company, USA) with a 48-channel head coil. Before the experiment, a T1-weighted image (MPRAGE) was collected as high-resolution anatomical reference (voxel size = 1×1×1mm^3^, repetition time (TR) = 2189.12ms, echo time (TE) = 2.96ms, flip angle = 8°, field of view (FoV) 256 mm, 156 slices). T2*-weighted gradient-echo echo-planar images were collected for the localizer experiment (TR = 1750ms, TE = 30ms, flip angle = 75°, FoV = 220mm × 220mm^2^, resolution = 2.6 × 2.6 mm^3^, 58 interleaved ascending slices, slice thickness = 2.6mm) and the decoding experiment (TR = 1750ms, TE = 35ms, flip angle = 75°, FoV = 220mm × 220mm^2^, resolution = 2.3 × 2.3 mm^3^, 52 interleaved ascending slices, slice thickness = 2.3mm) respectively.

MRI data were converted and organized into ‘Brain Imaging Data Structure’ (BIDS) format using dcm2bids (Boré et al., 2023) to ensure standardization and anonymization of data (Gorgolewski et al., 2016). Data were preprocessed with the preprocessing pipeline fMRIPrep 23.1.4 (Esteban et al., 2019) with the default processing steps incorporating the software packages: FSL, Freesurfer, ANTs and AFNI. First, each T1-weighted volume was corrected for intensity nonuniformity and skull-stripped to reconstruct brain surfaces. Brain-extracted T1-weighted images were then spatially normalized to the MNI152Lin2009cAsym standard space. Brain tissue segmentation of CSF, white matter, and gray matter was performed on the brain-extracted T1-weighted image. Functional data were slice time-corrected, motion-corrected and coregistered to the normalized T1-weighted template (for further details, including software packages for each preprocessing step in FMRIPrep, see the online documentation under https://fmriprep.org/en/stable/). Finally, we applied spatial smoothing using a 3D Gaussian kernel (univariate analysis, FWHM= 6 mm; MVPA analysis, FWHM = 2mm) using SPM 12 (Wellcome Department of Imaging Neuroscience, London, United Kingdom).

Numerosity processing has been shown to interact with spontanenous eye movement (Knops et al., 2009). To minimize the potential confounds of eye movement on brain activation, we computed eye displacement and included it in the Generalized Linear Model (GLM) as a nuisance regressor. Eye movement data were acquired using BIDSMReye (Gau et al., 2024), which is bidsified based on DeepMReye to decode eye motion from fMRI time series data, in particular from the MR signal of the eyeballs (Frey et al., 2021). Each run, the following values were computed: variance for the X gaze position, variance for the Y gaze position, framewise gaze displacement, number of X gaze position outliers, number of Y gaze position outliers, and number of gaze displacement outliers. Outliers were robustly estimated using an implementation of the median rule (Carling, 2000).

## Univariate analysis

The univariate fMRI analysis was conducted using bidspm (v3.0.0 by Gau et al.,) and SPM12 (version 7771) on MATLAB (R2019b). The pre-processed images for each participant were analyzed using a general linear model (GLM). Motion parameters (three translational motion, three rotational motion parameters), eye displacement (extracted from DeepMReye; Gau et al., 2024) and button presses were modelled as regressors of no interests. The contrast images for each condition against baseline were computed for each subject individually. The individual contrasts were further smoothed with a 6-mm FWHM kernel and entered into a random effects model for the second-level analysis. P-values were set at p < 0.05 FWE corrected, with probabilistic approach for threshold-free cluster enhancement (pTFCE) (Figure S1).

## Multivariate pattern analysis

For the numerosity decoding task, a general linear model was implemented for each subject to estimate beta maps associated with each numerosity (2,3,4,5) separately for the five stimulus conditions. For each level of numerosity within each condition, two subsets with different low-level features were modelled together as one beta to control for the low-level features of the stimuli. Motion parameters (three translational motion, three rotational motion parameters), eye displacement and button presses were modelled as regressors of no interests. For each run, one beta estimate is extracted for each 20 regressors of interest for further MVPA.

The MVPA analysis was performed in CoSMoMVPA (Oosterhof et al., 2016, http://www.cosmomvpa.org/), implemented in MATLAB. Classification analyses were performed using a multiclasses linear support vector machine (SVM) classifiers as implemented in LIBSVM (Chang & Lin, 2011). To explore the brain regions that can decode numerosity within each stimulus condition, we implemented a whole-brain searchlight approach to investigate which regions across the entire cortex could decode numerosity. A sphere of 3-voxels radius moved across the whole brain, where in each step, each voxel became the center of the searchlight sphere. Within each sphere, cross-modal decoding was implemented in the same manner described in the ROI analysis. The searchlight analysis results in a whole brain cross-modal decoding accuracy map for each subject. The individual accuracy maps were smoothed with 8 mm FWHM smoothing kernel and entered in a second-level model. A one-sample t test was performed to test if the cross-modal decoding accuracy was above the chance level (25%). P values were set at p < 0.001 for cluster-level correction corrected for multiple comparisons using pTFCE (probabilistic approach for threshold-free cluster enhancement). For the numerosity condition that could be decoded, a conjunction analysis was conducted on the accuracy maps from the different conditions, using a threshold of p< 0.001 (uncorrected), to identify regions in which numerosity could be decoded across all conditions. The peak coordinates from conjunction analysis were then used along with activation map to create ROIs for further the multivariate analysis. To define each region of interest (ROI), a 10 mm-radius sphere was first centered on each of the peak coordinate. This sphere was then progressively expanded and intersected with the activation map until the ROI reached approximately 300 voxels. When a sufficient number of voxels was not available, the intersection step was omitted, and a simple sphere around 300 voxels was drawn instead.

To ensure that successful decoding was not driven by low-level features specific to one subset of stimuli, we conducted a control analysis in which a classifier was trained on data from both subsets but tested only on one subset. If decoding relied on discriminative low-level features from one subset, it should fail when tested on the other subset. We reported classifier performance as the average across the two subsets used as the testing set. This analysis was performed for all three non-symbolic conditions.

To assess the existence of the distance effect in regions that identified through conjunction analysis (Dehaene et al., 1990; Nieder et al., 2002), we performed pairwise decoding between different numerosities within each condition, then averaged the decoding accuracy based on numerical distance, resulting in three groups. The first group corresponds to a numerical difference of one, the second group to a difference of two, and the third group to a difference of three. A linear regression model was then applied to the decoding accuracy across these groups. A linear increase in the decoding accuracy across groups would indicate the presence of a distance effect. To minimize multiple comparisons; ROIs were grouped into three cortical regions: occipital, parietal and frontal ROI regions. False discovery rate (FDR) was applied to P values across regions.

Cross-classification was performed in the ROIs that allowed to successfully decode numerosities to further test whether a shared numerosity representation existed across different conditions in these regions. Before the cross-classification, the pattern of each numerosity within each condition was demeaned individually to ensure the mean activity across conditions was mathematically equated. For each pair of conditions, a multiclass linear SVM classifier (C parameter = 1) was trained on one condition and tested on the other condition to classify four numerosities for each subject individually, and vice versa. The classifier was trained using a cross-validation leave-one-out procedure, with n-1 runs used for training and the remaining one run used for testing. We implemented an ANOVA-based feature selection on the training data within each cross-validation fold to use the most informative 150 voxels in each subject and each ROI. The normalization parameters from the training data were applied to the test data, and the performance of the SVM classifier was evaluated on the test data (1 run). The previous step was repeated 20 times (N-fold cross-validation) where in each fold the classifier was tested on a different run. A single classification accuracy value was obtained by averaging the classification accuracies for all the cross-validation folds. Successful performance above the chance level (25%) would indicate that specific ROIs are capable of decoding numerosities within the condition.

## Statistical significance

Statistical significance in the multivariate classification analyses was assessed using nonparametric tests permuting condition labels and bootstrapping (Stelzer et al., 2013). Each permutation step included shuffling of the condition labels and rerunning the classification 100 times on the single-subject level. Next, we applied a bootstrapping procedure to obtain a group-level null distribution that is representative of the whole group. From each individual’s null distribution, one value was randomly chosen and averaged across all of the participants. This step was repeated 10,000 times, resulting in a group-level null distribution of 10,000 values. The statistical significance of the MVPA results was estimated by comparing the observed result to the group-level null distribution. This was done by calculating the proportion of observations in the null distribution that had a classification accuracy higher than the one obtained in the real test. To account for the multiple comparisons, all p values were corrected using FDR correction method.

## Representational similarity analysis (RSA)

To further explore the differences in the representational formats used to encode numerosities, we relied on RSA. RSA allow capturing the (dis)similarity structure of numerosities across different representational formats and sensory modalities by testing which theoretical model best explains the observed data. To remove spatially correlated noise and yields generally more reliable dissimilarity estimates, multivariate noise normalization (Walther et al., 2016) was applied to the beta coefficients obtained for each stimulus by using The Decoding Toolbox (Hebart et al., 2015). For each ROIs that allowed to successfully decode numerosities, individual neural representational dissimilarity matrices (RDMs) were constructed using 1-Pearson correlation, characterized as 1 minus the Pearson correlation of the β-weight patterns across all pairwise conditions.

We built three theoretical models that we thought could represent how brain regions encode our different numerosities (see Figure 1C). First, the Condition-Numerosity (CondNum) model assumes that numerosities are more similar within than across each condition. The dissimilarity between numerosity from different conditions was arbitrarily set to 1 (maximum dissimilarity), while within condition dissimilarity is based on numerical distance. For example, the dissimilarity of numerosity 2 compared to numerosities 3, 4, and 5 increases progressively, with values of 0.16, 0.33, and 0.5 respectively. Second, the Modality-Numerosity model (ModalNum) assumes that numerosities from the same modality (e.g. vision) are more similar than those from different modalities even across format. The dissimilarity between numerosities from different modalities is set to 1, while within the same condition, dissimilarity is determined by their numerical distance following the same logic as the CondNum model. Third, the Format-Numersity model (FormatNum) follows a similar principle with the previous model, but focuses on presentation formats (e.g., sequential vs. simultaneous). Numerosities in the same format are more similar than those from different formats, with dissimilarities between formats set to 1. Within a format, numerical distance determines the dissimilarity, with values of 0.16, 0.33, and 0.5 for numerosities 2, 3, 4, and 5; respectively.

To assess the between-subjects reliability of the neural data, we computed the noise ceiling of the data for each fROI. The bottom half of RDMs was first converted into a vector and z-transformed. A lower bound was computed by the following approach: the vector of one subject was left out and those of remaining subjects were averaged and correlated with the left-out subject’s vector. The same process was repeated for all subjects, and the resulting correlation coefficients were averaged. An upper bound was computed by correlating between each participant’s individual RDM and the group-average RDM (Nili et al., 2014).

We correlated the individual neural RDM with each theoretical model to assess which model best describes the neural data. The median correlations across subjects were tested for significance with Wilcoxon signed-rank tests. Results were corrected for multiple comparisons using FDR correction.

## Partial correlations and variance partitioning

To determine the contribution of each individual model when considered in conjunction with the other models, we performed two additional analyses: partial correlations, in which each model was correlated (Pearson r) while partialling out the other two models, and variance partitioning based on multiple linear regression (see Groen et al., 2018; Ritchie et al., 2021 for similar approach). In the present analysis, CondNum (*a*), ModalNum (*b*) and FormatNum (*c*) models were converted into vectors and included as predictors in a set of multiple linear regression analyses. Each model was tested individually, as well as in all possible pairwise and three-way combinations, to predict individual participants’ RDMs. Subsequently, seven R^2^ values were obtained: R^2^, R^2^, R^2^, R^2^, R^2^, R^2^_y•bc_, R^2^. The first six R^2^ were based on either individual models or combinations of pairwise model, while the last R^2^ indicated the full model. By comparing the explained variance (R^2^) of full model to the R^2^ obtained when each model is left out one at a time, we can infer the amount of variance uniquely explained by the excluded model. The results were visualized in a Euler diagram using EulerAPE software (Micallef & Rodgers, 2014).

Partial correlations were calculated for each individual participant separately. The median correlations across subjects were tested for significance with Wilcoxon signed-rank tests. Results were corrected for multiple comparisons using FDR correction.

## Acknowledgements

Computational resources have been provided by the supercomputing facilities of the Université catholique de Louvain (CISM/UCL) and the Consortium des Équipements de Calcul Intensif en Fédération Wallonie Bruxelles (CÉCI) funded by the Fond de la Recherche Scientifique de Belgique (F.R.S.-FNRS) under convention 2.5020.11 and by the Walloon Region. Authors thank Michael Andres and Marie-Pascale Noel for their advises on the design development.

## Funding

The project was funded in parts by the Belgian Excellence of Science (EOS) program (Project No. 30991544) awarded to O.C. O.C. is a senior research associate at the Fond National de la Recherche Scientifique de Belgique (FRS-FNRS). Y.Y. is funded by the Chinese Scholarship Council.

## Conflict of interest disclosure

None.

## Author Contributions

Y.Y. and O.C. designed research. Y.Y., I.S., A.A and F.C. collected data. Y.Y. analyzed data under the supervision of OC. I.S. and R.G. provided support on data analysis scripts and toolbox. Y.Y. M.F., I.T. and O.C. wrote the manuscript. All authors reviewed and edited the manuscript. Y.Y. and O.C. acquired funding.

## Data availability statement

Raw f/MRI data are not publicly available as full anonymity of the participants cannot be guaranteed, even after defacing the MRI images and due to the lack of explicit consent from our participants. The results of searchlight map, ROI cross-classification and RSA can be found on OSF (https://osf.io/x9awu/?view_only=548ffa994aaf44e8bb3a3dbd8acf15e6).

---

**Figure S1.**
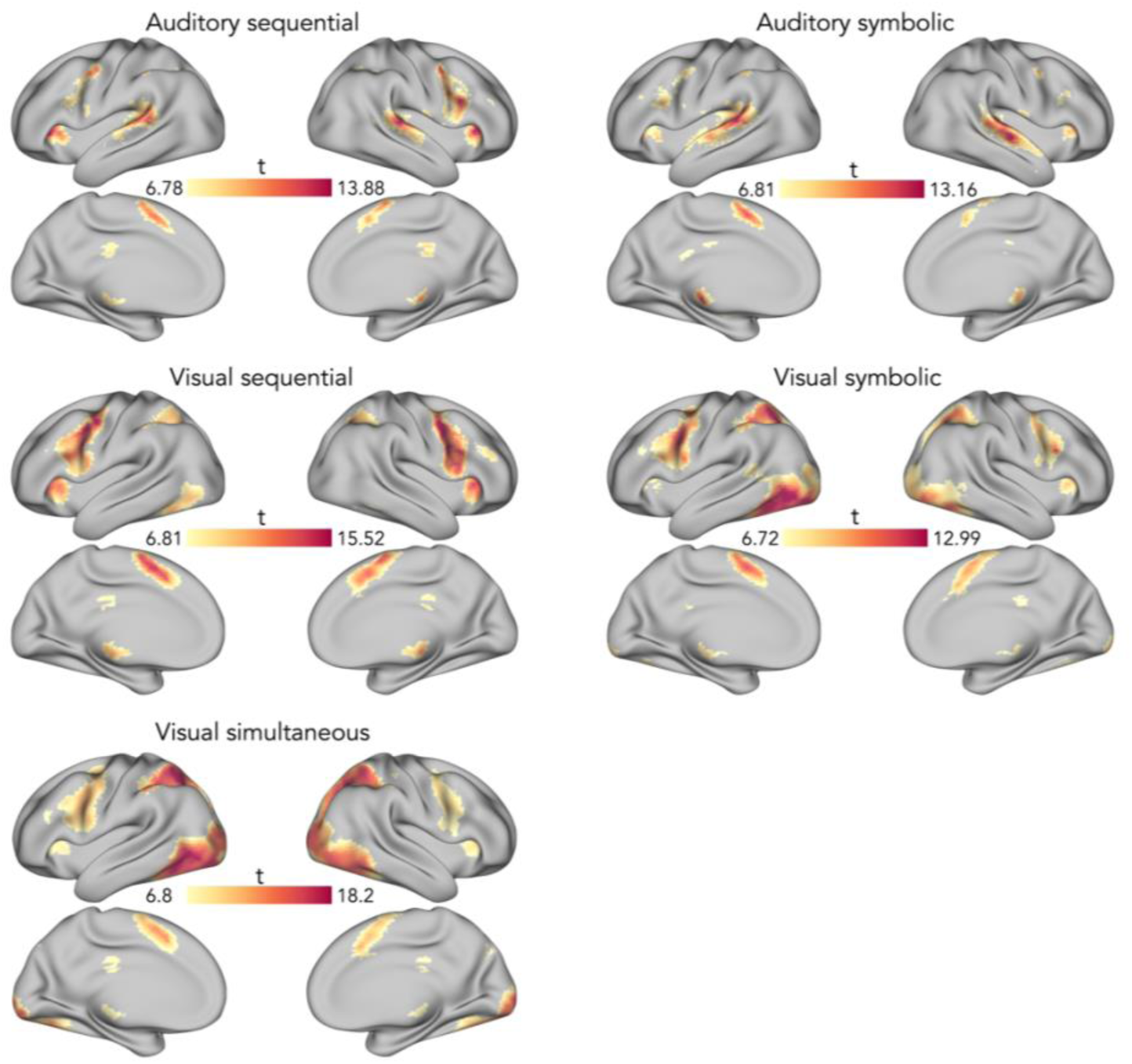
The univariate results of the contrast each stimuli condition against baseline. P-values were set at p < 0.05 for cluster-level correction controlling family-wise error (FWE) with probabilistic threshold-free cluster enhancement (pTFCE). Results are displayed on an inflated brain (Marcus et al., 2013).

**Figure S2.**
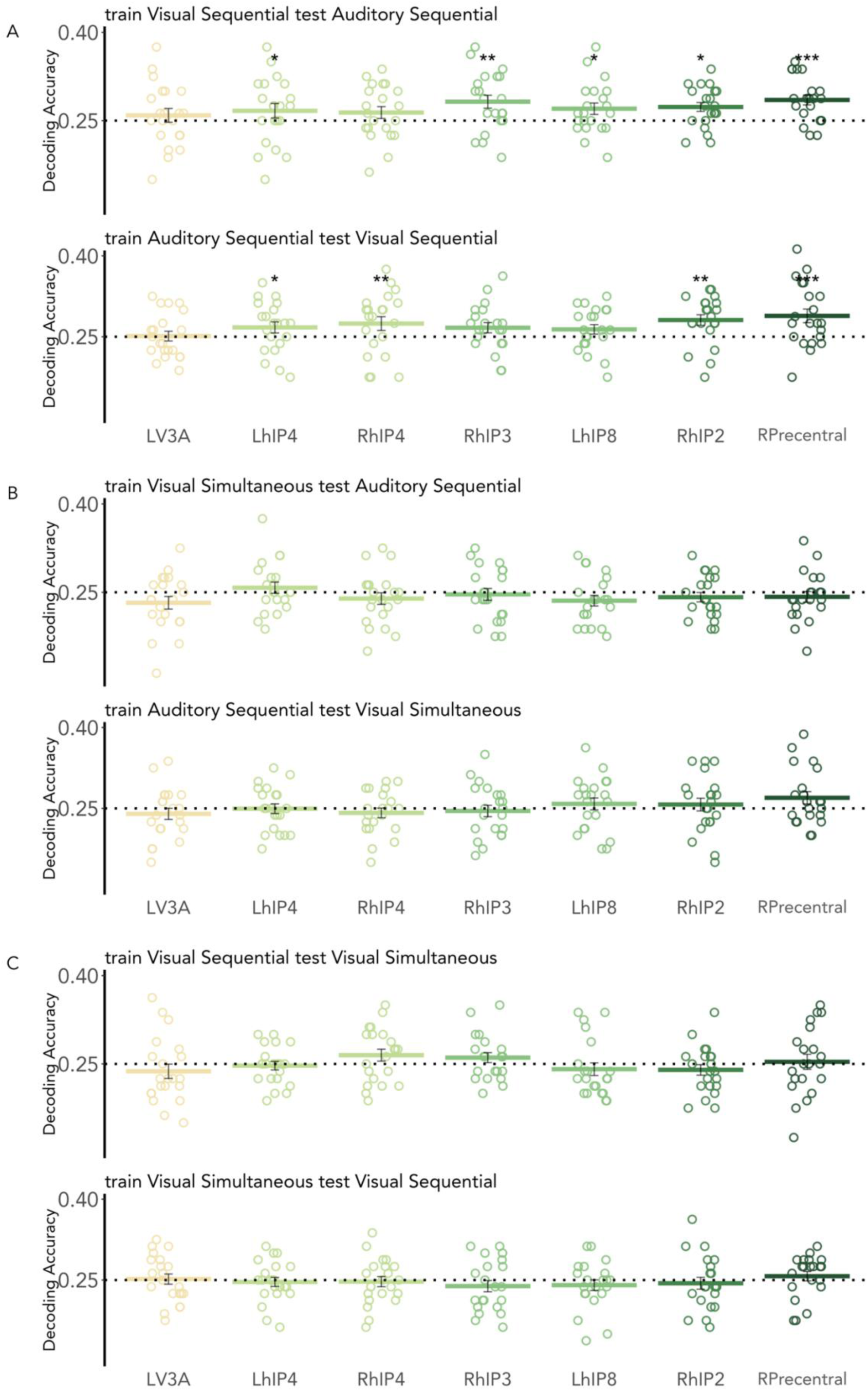
Cross-classification results for each classification direction within every pair of non-symbolic numerosity conditions. Chance level (dotted line) is 25%. Cross-bar colors encode the ROIs, shifting from light to dark green as the regions move along the cortical surface from posterior to anterior. Individual points reflect individual subjects’ results. Results are FDR corrected (*p < 0.05, **p < 0.01, ***p < 0.001). Error bars represent standard error of mean. **A.** Visual sequential v.s. auditory sequential numerosities. **B.** Auditory sequential v.s. Visual simultaneous numerosities. **C.** Visual sequential v.s. Visual simultaneous numerosities.

**Figure S3.**
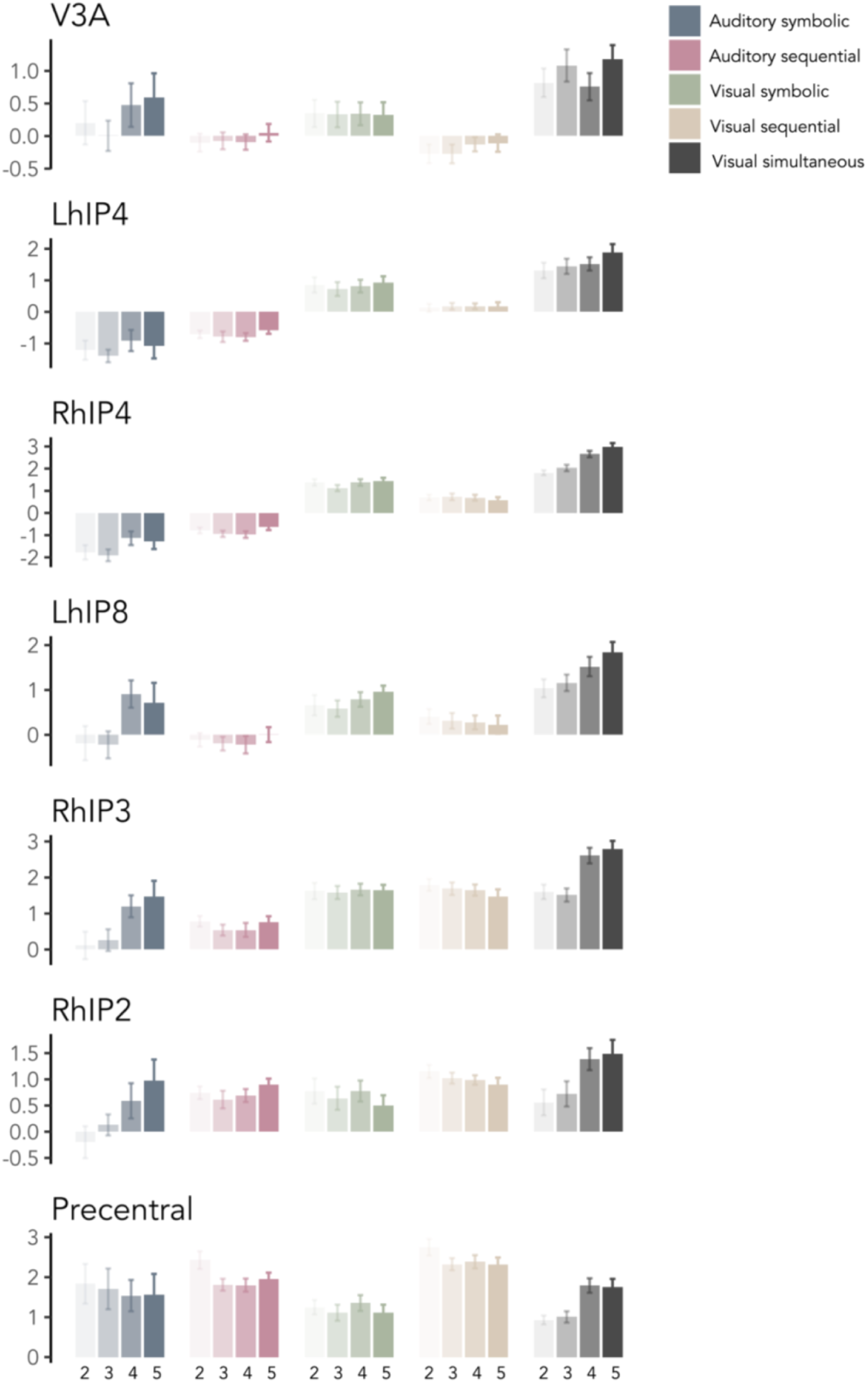
Beta parameter estimates extracted from the ROIs defined by conjunction map. Error bars represent the standard error of the mean. Increasing bar alpha levels represent increasing numerosity.

**Figure S4.**
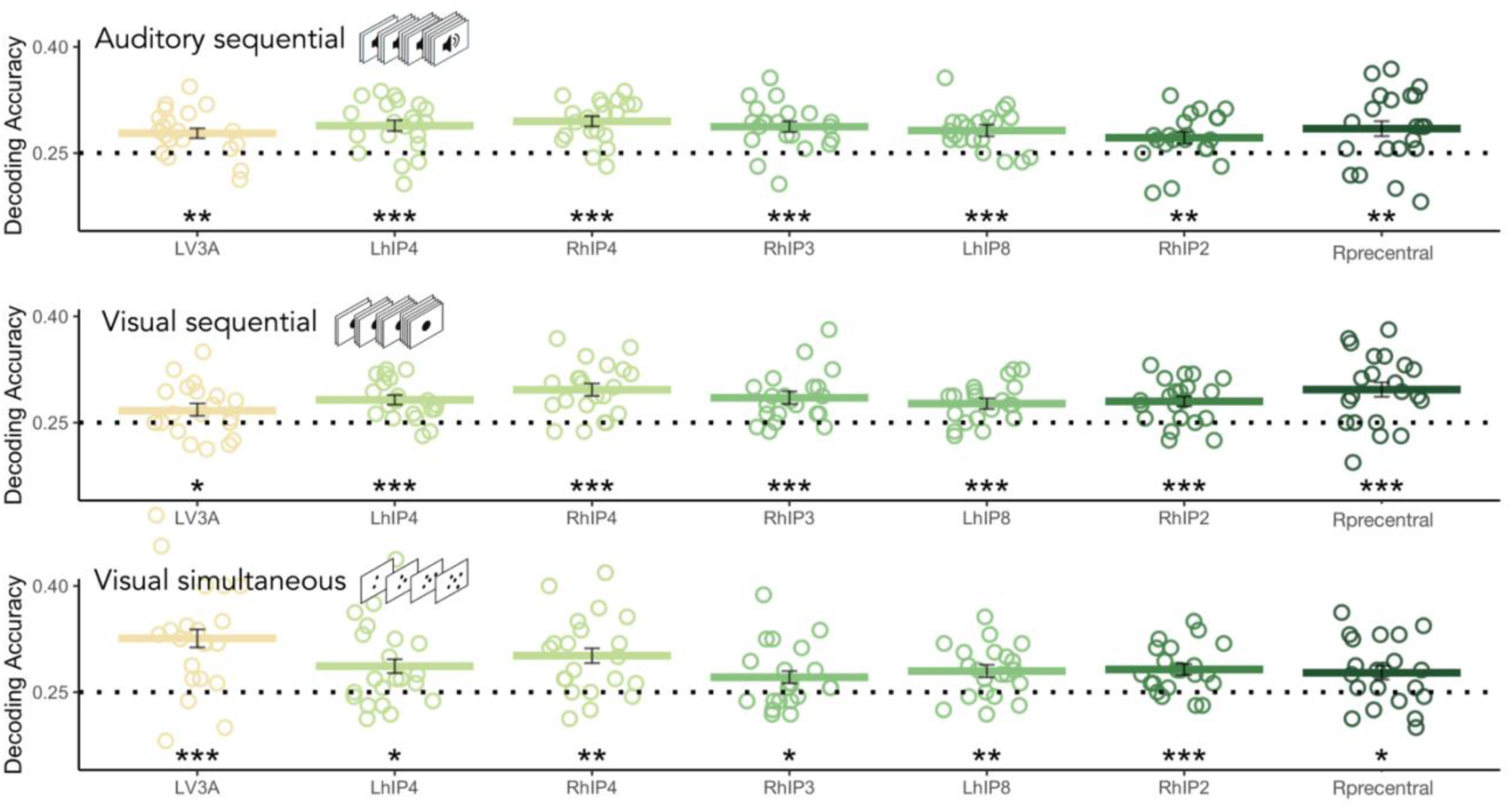
Decoding accuracy when the classifier was trained on data from both stimulus subsets but tested on only one subset. Classifier performance reported here is the average across the two subsets used as the testing set. Results are FDR corrected (*p < 0.05, **p < 0.01, ***p < 0.001). Error bars represent standard error of mean.

## Supplementary Tables and Figures

**Table S1.** Detailed information for dots arrays for each subsets of each numerosity. Convex hull: the area of the smallest convex polygon containing all the elements. Inter-distance: the average of the distances between all the possible pairs of elements. Density: the number of elements (n) divided by the total occupied area (D = n/CH). Radius: the specific dimension of a single dot. Total perimeter: the sum of the perimeters (contours) of all elements. Total area: the sum of the areas (surfaces) of all elements (Zanon et al., 2022). For each numerosity in each condition, two dots arrays are generated within each set and are randomly assigned across participants.

|  |  | convex hull | density(N/convex hull) | inter-distance | total area | total perimeter | radius |
| --- | --- | --- | --- | --- | --- | --- | --- |
| 2 | subset1 | 90,0019947 | 0,02222173 | 13,8721102 | 40 | 31,7066184 | 2,52313252 |
|  |  | 90,0012929 | 0,0222219 | 13,8719721 | 40 | 31,7066184 | 2,52313252 |
| 3 | subset1 | 90,0002661 | 0,03333323 | 8,3532217 | 40 | 38,8325183 | 2,06012908 |
|  |  | 90,0000771 | 0,0333333 | 8,254024 | 40 | 38,8325183 | 2,06012908 |
| 4 | subset1 | 90,0018426 | 0,04444354 | 6,92706994 | 40 | 44,8399297 | 1,78412412 |
|  |  | 90,0032252 | 0,04444285 | 7,82048802 | 40 | 44,8399297 | 1,78412412 |
| 5 | subset1 | 90,0008139 | 0,05555505 | 7,23772541 | 40 | 50,1325655 | 1,59576912 |
|  |  | 90,0021445 | 0,05555423 | 6,62875581 | 40 | 50,1325655 | 1,59576912 |
|  |  | convex hull | density(N/convex hull) | inter-distance | total area | total perimeter | radius |
| 2 | subset2 | 42,5794864 | 0,04697098 | 7,99940581 | 23,404237 | 24,2530953 | 1,93 |
|  |  | 42,5673277 | 0,04698439 | 7,99625244 | 23,404237 | 24,2530953 | 1,93 |
| 3 | subset2 | 67,8586757 | 0,04420953 | 7,9988023 | 35,1063554 | 36,3796429 | 1,93 |
|  |  | 78,941948 | 0,03800261 | 7,99661894 | 35,1063554 | 36,3796429 | 1,93 |
| 4 | subset2 | 108,840337 | 0,03675108 | 8,00278454 | 46,8084739 | 48,5061906 | 1,93 |
|  |  | 110,96689 | 0,03604679 | 7,99735734 | 46,8084739 | 48,5061906 | 1,93 |
| 5 | subset2 | 136,016481 | 0,03676025 | 7,99581817 | 58,5105924 | 60,6327382 | 1,93 |
|  |  | 124,809709 | 0,04006099 | 8,00062427 | 58,5105924 | 60,6327382 | 1,93 |

**Table S2.** Timing information (mean (SD)) for visual and auditory sequential numerosity conditions. Dots/Beeps duration represents the duration of individual dots/beeps within a sequence. Inter dot/beep interval represents the duration between individual dots/beeps. Total sound/beep duration represents the summed duration of dots/beeps in a sequence, excluding the intervals. Total length is the combined duration of the total dot/beep duration and the total inter-dot/beep intervals. For each numerosity in each condition, two sequences are generated within each set and are randomly assigned across participants.

| visual sequential | Dots Duration |  | Inter Dot Interval |  | Total Sound Duration |  | Total Length |  |
| --- | --- | --- | --- | --- | --- | --- | --- | --- |
| numerosity | set1 | set2 | set1 | set2 | set1 | set2 | set1 | set2 |
| 2 | 0.634(0.254) | 0.240(0.035) | 0.653(0) | 0.180(0) | 1.267 | 0.48 | 1.92 | 0.66 |
| 3 | 0.422(0.121) | 0.240(0.027) | 0.326(0.524) | 0.180(0.088) | 1.267 | 0.720 | 1.92 | 1.08 |
| 4 | 0.317(0.067) | 0.240(0.036) | 0.217(0.045) | 0.180(0.089) | 1.267 | 0.960 | 1.92 | 1.5 |
| 5 | 0.240(0.020) | 0.240(0.020) | 0.180(0.095) | 0.180(0.095) | 1.200 | 1.200 | 1.92 | 1.92 |
| auditory sequential | Beeps Duration |  | Inter Dot Interval |  | Total Sound Duration |  | Total Length |  |
| numerosity | set1 | set2 | set1 | set2 | set1 | set2 | set1 | set2 |
| 2 | 0.528(0.168) | 0.187(0.032) | 0.544(0) | 0.175(0.035) | 1.056 | 0.375 | 1.6 | 0.55 |
| 3 | 0.352(0.079) | 0.200(0.032) | 0.272(0.009) | 0.150(0.058) | 1.056 | 0.6 | 1.6 | 0.9 |
| 4 | 0.264(0.041) | 0.200(0.019) | 0.181(0.072) | 0.150(0.071) | 1.056 | 0.8 | 1.6 | 1.25 |
| 5 | 0.211(0.058) | 0.200(0.020) | 0.136(0.044) | 0.150(0.080) | 1.056 | 1 | 1.6 | 1.6 |

**Table S3.** Clusters identified in conjunction analysis of three non-symbolic searchlight accuracy map. Results are corrected for multiple comparison using FWE correction at the cluster level. Coordinates are reported in MNI space.

| Region | Cluster-level p<br>(FWE-corr) | Peak_level<br>t | x | y | z (mm) |
| --- | --- | --- | --- | --- | --- |
| Left hIP8 | 0.002 | 5.90 | -22 | -72 | 48 |
|  |  | 4.93 | -19 | -74 | 34 |
|  |  | 4.76 | -24 | -65 | 39 |
| Left V3A | <0.001 | 5.46 | -19 | -88 | 13 |
| Left V1 |  | 5.36 | 2 | -79 | 25 |
| Left hIP4 |  | 5.31 | -33 | -79 | 20 |
| Right hIP2 | 0.002 | 5.44 | 53 | -32 | 41 |
|  |  | 4.90 | 44 | -41 | 53 |
| Right hIP3 | <0.001 | 5.40 | 35 | -58 | 50 |
| Right hIP4 |  | 5.18 | 35 | -76 | 27 |
| Right hIP7 |  | 5.04 | 30 | -81 | 23 |
| Precentral | 0.012 | 5.31 | 49 | 10 | 27 |
|  |  | 4.18 | 44 | 6 | 39 |

**Table S4.**
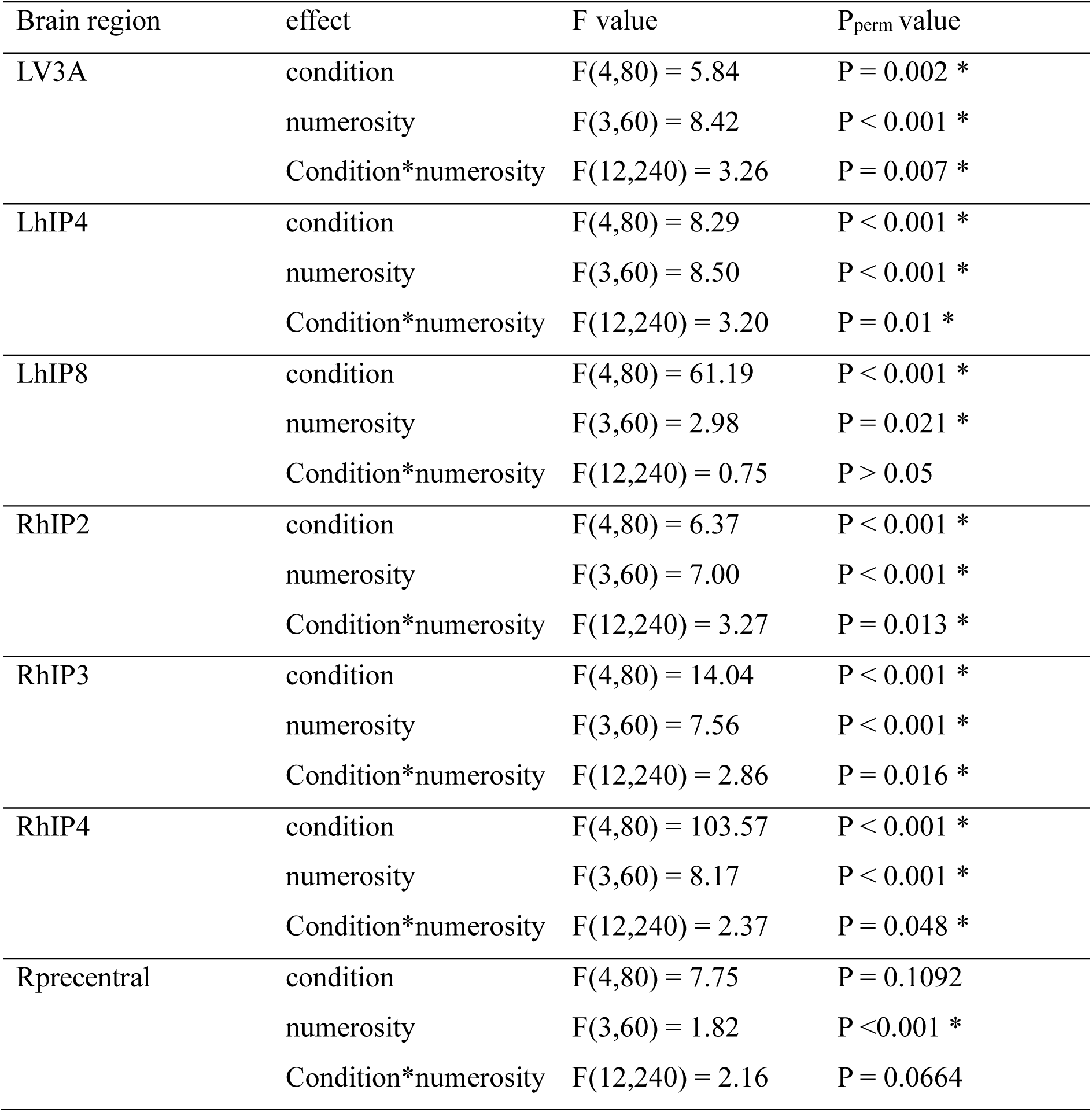
Statistical results of the non-parametric ANOVA for beta estimates parameters.

**Table S5.**
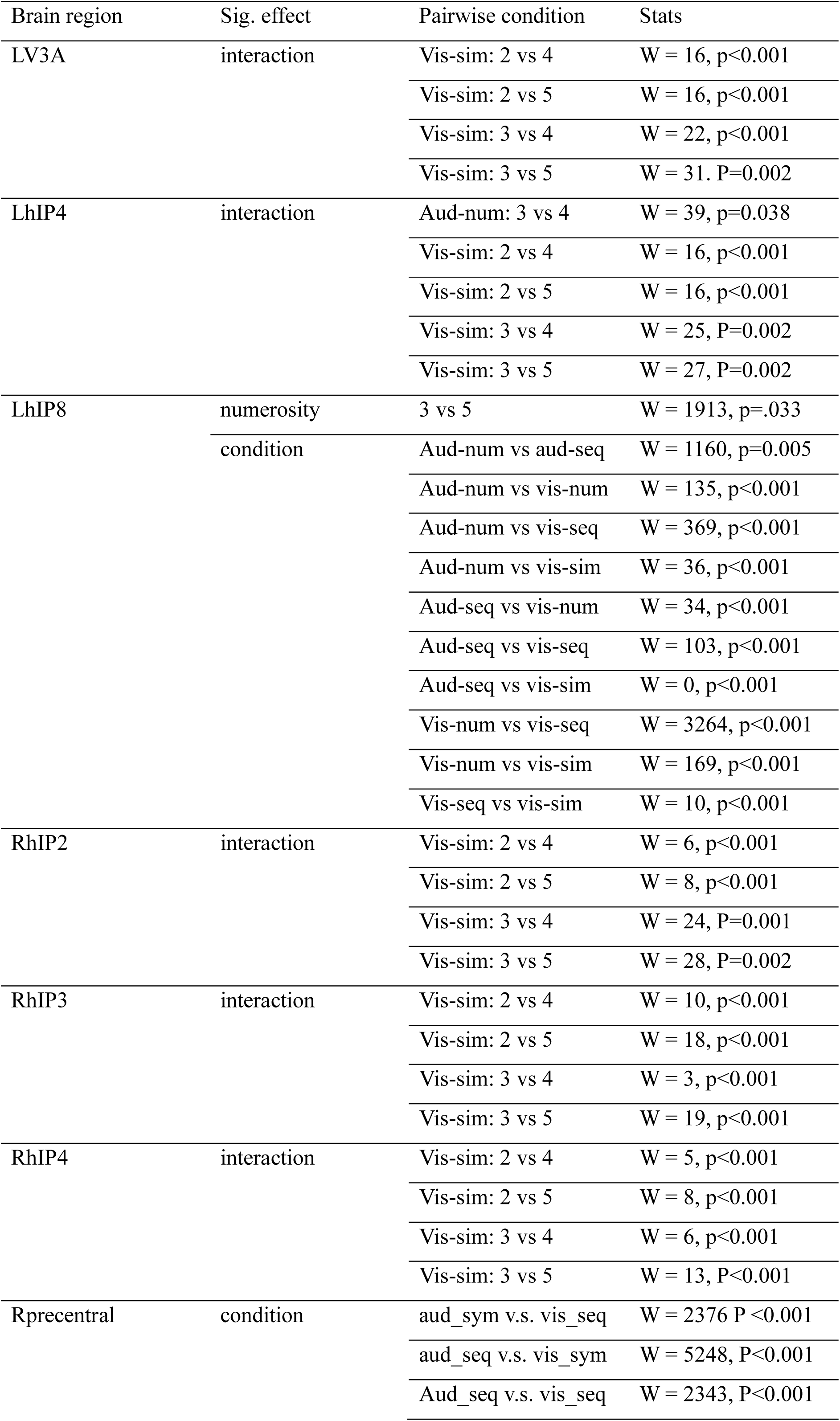
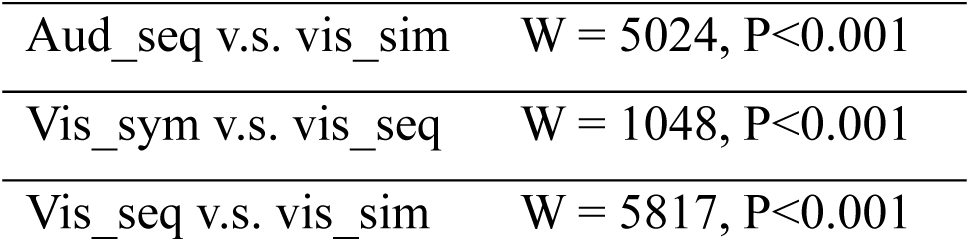
Post hoc Wilcoxon pairwise comparisons following significant main effects or interactions in beta estimates parameters.

| Brain region | Sig. effect | Pairwise condition | Stats |
| --- | --- | --- | --- |
| LV3A | interaction | Vis-sim: 2 vs 4 | W = 16, p<0.001 |
|  |  | Vis-sim: 2 vs 5 | W = 16, p<0.001 |
|  |  | Vis-sim: 3 vs 4 | W = 22, p<0.001 |
|  |  | Vis-sim: 3 vs 5 | W = 31, P=0.002 |
| LhIP4 | interaction | Aud-num: 3 vs 4 | W = 39, p=0.038 |
|  |  | Vis-sim: 2 vs 4 | W = 16, p<0.001 |
|  |  | Vis-sim: 2 vs 5 | W = 16, p<0.001 |
|  |  | Vis-sim: 3 vs 4 | W = 25, P=0.002 |
|  |  | Vis-sim: 3 vs 5 | W = 27, P=0.002 |
| LhIP8 | numerosity | 3 vs 5 | W = 1913, p=.033 |
|  | condition | Aud-num vs aud-seq | W = 1160, p=0.005 |
|  |  | Aud-num vs vis-num | W = 135, p<0.001 |
|  |  | Aud-num vs vis-seq | W = 369, p<0.001 |
|  |  | Aud-num vs vis-sim | W = 36, p<0.001 |
|  |  | Aud-seq vs vis-num | W = 34, p<0.001 |
|  |  | Aud-seq vs vis-seq | W = 103, p<0.001 |
|  |  | Aud-seq vs vis-sim | W = 0, p<0.001 |
|  |  | Vis-num vs vis-seq | W = 3264, p<0.001 |
|  |  | Vis-num vs vis-sim | W = 169, p<0.001 |
|  |  | Vis-seq vs vis-sim | W = 10, p<0.001 |
| RhIP2 | interaction | Vis-sim: 2 vs 4 | W = 6, p<0.001 |
|  |  | Vis-sim: 2 vs 5 | W = 8, p<0.001 |
|  |  | Vis-sim: 3 vs 4 | W = 24, P=0.001 |
|  |  | Vis-sim: 3 vs 5 | W = 28, P=0.002 |
| RhIP3 | interaction | Vis-sim: 2 vs 4 | W = 10, p<0.001 |
|  |  | Vis-sim: 2 vs 5 | W = 18, p<0.001 |
|  |  | Vis-sim: 3 vs 4 | W = 3, p<0.001 |
|  |  | Vis-sim: 3 vs 5 | W = 19, p<0.001 |
| RhIP4 | interaction | Vis-sim: 2 vs 4 | W = 5, p<0.001 |
|  |  | Vis-sim: 2 vs 5 | W = 8, p<0.001 |
|  |  | Vis-sim: 3 vs 4 | W = 6, p<0.001 |
|  |  | Vis-sim: 3 vs 5 | W = 13, P<0.001 |
| Rprecentral | condition | aud_sym v.s. vis_seq | W = 2376 P <0.001 |
|  |  | aud_seq v.s. vis_sym | W = 5248, P<0.001 |
|  |  | Aud_seq v.s. vis_seq | W = 2343, P<0.001 |

|  |  |
| --- | --- |
| Aud_seq v.s. vis_sim | W = 5024, P<0.001 |
| Vis_sym v.s. vis_seq | W = 1048, P<0.001 |
| Vis_seq v.s. vis_sim | W = 5817, P<0.001 |

## Statistical results of correlation between neural dissimilarity matrix and theoretical models

We tested the correlation between the neural dissimilarity matrix (DSM) of each ROI in each subject with three theoretical models (Figure 5A, 5B). In the early visual cortex, the median correlations of the three model RDMs were all significant [all: W(21) < 7, r_bc_ < 0.0303, p ≤ 0.001]. Additionally, we also observed significant differences between the condition numerosity model (CondNum) versus the format numerosity model (FmtNum), and the format numerosity model versus the modality numerosity model (ModalNum) [W(21) < 56, r_bc_ < 0.2424, p ≤ 0.003]. In the posterior parietal subregion left hIP4, the median correlations of the three model RDMs were all significant [all: W(21) = 0, r_bc_ = 0, p ≤ 0.001]. The differences between every two models were also significant [all: W (21) ≤ 64, r_bc_ ≤ 0.2771, p ≤ 0.0117]. In the posterior parietal subregions right hIP4, the median correlations of the three model RDMs were all significant [all: W(21) = 0, r_bc_ = 0, p ≤ 0.001]. The differences between every two models were also significant [all: W (21) ≤ 59, r_bc_ ≤ 0.2554, p ≤ 0.0015]. In the lateral parietal left hIP8 region, all three models showed significant partial correlations [all: W(21) ≤ 4, r_bc_ ≤ 0.0173, p ≤ 0.001]. Additionally, there was a significant difference between the condition numerosity model versus the modality numerosity model [W(21) = 65, r_bc_ = 0.2814, p = 0.0078]. In the lateral parietal right hIP3 region, all three models showed significant partial correlations [all: W(21) = 0, r_bc_ = 0, p ≤ 0.001]. In the anterior parietal subregions left hIP2, all three models showed significant partial correlations [all: W(21) ≤ 6, r_bc_ ≤ 0.026, p ≤ 0.001]. There was also a significant difference between condition numerosity model versus format numerosity model [W(21) = 8, r_bc_ = 0.0346, p = 0.0012]. In the frontal Precentral region, all three models showed significant partial correlations [all: W(21) ≤ 21, r_bc_ ≤ 0.0909, p ≤ 0.001]. There are significant difference between condition numerosity model versus modality numerosity model [W(21) = 49, r_bc_ = 0.2121, p = 0.0057].

## Statistical results of partial correlation between neural dissimilarity matrix and theoretical models

To assess the independent contribution of each model to the neural activity, we computed partial correlations between the models and neural activities. In the early visual cortex area V3A, significant partial correlations were found for all three models [all: W(21) ≤ 64, rbc ≤ 0.2771, p ≤ 0.012]. In the posterior parietal subregions left hIP4, significant partial correlations were observed for both the modality and format models [W (21) ≤ 11, rbc ≤ 0.0476, p ≤ 0.001]. In the posterior parietal subregions right hIP4, significant partial correlations were found for all three models [all: W (21) ≤ 49, rbc ≤ 0.2121, p ≤ 0.019]. In the lateral parietal left hIP8 region, all three models showed significant partial correlations [all: W(21) ≤ 71, rbc ≤ 0.3074, p ≤ 0.0032]. In the lateral parietal right hIP3 region, significant partial correlations were obtained for the modality and format model [both: W(21) = 0, rbc = 0, p ≤ 0.001]. In the anterior parietal subregions left hIP2, significant partial correlations were obtained for all three models [all: W(21) ≤ 9, rbc ≤ 0.0390, p ≤ 0.001]. Likewise, in the frontal Precentral region, all three models showed significant partial correlations [all: W(21) ≤ 51, rbc ≤ 0.2208, p ≤ 0.001].

## Beta parameter estimates

To examine the activity levels associated with different conditions and numerosities, and to determine whether specific conditions or numerosities consistently evoked higher or lower brain activity, we extracted univariate beta parameter estimates in each fROI where we observed successful numerosity decoding. We conducted non-parametric ANOVAs. In case of significant main effect or interaction effect, post hoc tests were conducted using a series of Wilcoxon tests, and were adjusted for multiple comparisons using the false discovery rate (FDR) methods (Benjamini & Yekutieli, 2001).

Six out of seven ROIs demonstrated significant main effects of numerosity, condition, as well as a significant numerosity × condition interaction (Figure S3). Post hoc pairwise comparisons revealed that the interaction effect was mostly driven by the significant beta difference in the visual simultaneous condition. Detailed results of the non-parametric ANOVA and post hoc Wilcoxon pairwise tests are provided in Table S4-S5.

## Notes

### Competing Interest Statement

The authors have declared no competing interest.

### Summary of Updates

figure revised; Supplemental files updated, reformatting text

